# DNA methylation during development and regeneration of the annelid *Platynereis dumerilii*

**DOI:** 10.1101/2020.11.13.381673

**Authors:** Anabelle Planques, Pierre Kerner, Laure Ferry, Christoph Grunau, Eve Gazave, Michel Vervoort

## Abstract

**Background:** Methylation of cytosines in DNA (5mC methylation) is a major epigenetic modification that modulates gene expression and is important for embryonic development and cell reprogramming in vertebrates. In mammals, 5mC methylation in promoter regions is linked to transcriptional repression. Transcription regulation by 5mC methylation notably involves the Nucleosome Remodeling and Deacetylase complex (NuRD complex) which bridges DNA methylation and histone modifications. Less is known about roles and mechanisms of 5mC methylation in non-vertebrate animals. In this paper, we study 5mC methylation in the marine annelid worm *Platynereis dumerilii,* an emerging evolutionary and developmental biology model capable of regenerating the posterior part of its body upon amputation. The regenerated region includes both differentiated structures and a growth zone consisting of stem cells required for the continuous growth of the worm.

**Results:** Using *in silico* and experimental approaches, we show that *P. dumerilii* displays a high level of DNA methylation comparable to that of mammalian somatic cells. 5mC methylation in *P. dumerilii* is dynamic along the life cycle of the animal and markedly decreases at the transition between larval to post-larval stages. We identify a full repertoire of mainly singlecopy genes encoding the machinery associated to 5mC methylation or members of the NuRD complex in *P. dumerilii* and show, through phylogenetic analyses, that this repertoire is close to the one inferred for the last common ancestor of bilaterians. These genes are dynamically expressed during *P. dumerilii* development, growth and regeneration. Treatment with the DNA hypomethylating agent Decitabine, impairs *P. dumerilii* larval development and regeneration, and has long-term effects on post-regenerative growth by affecting the functionality of stem cells of the growth zone.

**Conclusions:** Our data indicate high-level of 5mC methylation in the annelid *P. dumerilii,* highlighting that this feature is not specific to vertebrates in the bilaterian clade. Analysis of DNA methylation levels and machinery gene expression during development and regeneration, as well as the use of a chemical inhibitor of DNA methylation, suggest an involvement of 5mC methylation in *P. dumerilii* development, regeneration and stem cell-based post-regenerative growth. We also present data indicating that *P. dumerilii* constitutes a promising model to study biological roles and mechanisms of DNA methylation in non-vertebrate bilaterians and to provide new knowledge about evolution of the functions of this key epigenetic modification in bilaterian animals.

## BACKGROUND

Epigenetic modifications or marks refers to any transient chemical alterations of nucleic acids or histones, which do not modify the primary nucleic acid sequence and which can be transmitted from one generation of cells (and in some cases of organisms) to the next [1,2]. Epigenetic marks regulate gene expression and are therefore of paramount importance in most aspects of the biology of living organisms, including during development, regeneration and stem cells maintenance in animals [3,4].

DNA methylation is an important epigenetic modification, found in the three domains of life, and which has been the subject of intense study for many years [5–8]. In animals, DNA methylation mainly occurs through covalent addition of a methyl group on position 5 of a cytosine to form 5-methyl-cytosine (5mC). 5mC are mostly (or even exclusively in some species) found in cytosine-guanine dinucleotides, known as CpG sequences [9]. Abundance and distribution of 5mC strongly vary in different animal lineages. For example, in mammals, about 70-80% of CpGs throughout the genome are methylated in somatic tissue types. Unmethylated regions are largely restricted to dense clusters of CpGs, known as CpG islands (CGIs), which account for roughly two thirds of mammalian gene promoters. In the rare cases where CGI promoters are highly methylated, genes are stably transcriptionally repressed. In many non-vertebrates, methylated CpGs are mostly found within gene bodies [10,11]. The function(s) of this form of DNA methylation, which is referred to as “gene body methylation” (also found in vertebrates), is still largely unknown, but it has been hypothesized that it could be involved in homeostatic regulation of gene transcription [12].

DNA 5mC presence and roles rely on the activity of several classes of proteins that can be functionally classified according to their role: methylases that promote addition of methyl groups (“Writers”); proteins that oxidize 5mC and stimulate demethylation (“Modifiers”); and proteins that bind to methylated nucleotides, allowing interpretation of encoded information, for example in term of gene expression (“Readers”) [5,13,14]. Deposition of 5mC marks on DNA requires the action of evolutionary conserved DNA methyltransferases (Dnmts) [15]. In mammals, three families of Dnmts are found (Dnmt1, 2 and 3), each of which play specific roles [5,7]. Dnmt3 proteins are involved in the *de novo* addition of 5mC, while another member of the family, Dnmt1 is in charge of maintaining methylation pattern during replication. Dnmt1 function involves Uhrf1 (Ubiquitin Like with PHD And Ring Finger Domains 1 protein) which binds to both hemi-methylated DNA and Dnmt1, thereby recruiting Dnmt1 to methylated DNA sites [16]. Dnmt2 (also referred to as tRNA Aspartic Acid Methyltransferase 1) is a tRNA methylating enzyme seemingly not involved in DNA methylation, at least in mammals [17]. DNA demethylation, which results in the recovery of non-methylated cytosines, occurs either passively during cell division (in the absence of Dnmt1 function) or actively thanks to the Ten-Eleven Translocation (Tet) family enzymes and G/T Mismatch-Specific Thymine DNA Glycosylase (Tdg) proteins [18,19].

A series of proteins, known as methyl-CpG binding proteins (Mbp), recognize and bind methylated CpGs, acting as readout of DNA methylation by recruiting chromatin remodelers [20,21]. One key family of Mbp are methyl-CpG-binding domain (Mbd) proteins, which are found in many animals [22]. In mammals, seven members of the family are found and they mostly promote transcriptional silencing by interacting with a wide array of histone methylases and deacetylases [20,21]. One specific member, Mbd2, is part of the Nucleosome Remodeling and Deacetylase complex (NuRD complex), which bridges DNA methylation and histone modifications, and has been shown to regulate gene expression [23,24]. The NuRD complex is composed of several protein subunits: Mbd2 that binds methylated DNA, Chromodomain Helicase DNA Binding Proteins (Chd 3/4/5) which remodel chromatin, class I Histone Deacetylases (Hdac 1, 2, 3 and 8) that deacetylate histone tails and are associated to chromatin compaction and gene silencing, Retinoblastoma-Binding Protein (Rbbp4/7; also known as RbAp46/48) which is a histone chaperone, GATA Binding Protein (Gata2a/b), and Metastasis-Associated proteins (Mta1/2/3).

Most of what we know about DNA methylation in metazoans comes from studies conducted in a few model organisms, mainly mammals. While a handful of studies of DNA methylation in non-model organisms from various animal clades have been published recently *(e.g.,* [25–30]), little is known about the importance and roles of 5mC modifications in nonvertebrate species. We therefore decided to study DNA methylation in an emerging developmental biology model system, the marine annelid worm *Platynereis dumerilii.* Annelids belong, together with phyla such as molluscs and platyhelminthes, to lophotrochozoans, one of the three branches of bilaterian animals, distinct to those to which belong vertebrates (deuterostomes), and arthropods and nematodes (ecdysozoans) [31]. *P. dumerilii* has a complex life cycle composed of several phases [32] starting by a three-day long embryonic and larval development that gives rise to small larvae with three segments bearing appendages (parapodia). These larvae then metamorphose into small juvenile worms that enter a long phase of juvenile growth during which they add additional segments one by one in the posterior part of their body (a process known as posterior growth or elongation) [33]. Posterior growth relies on the presence of a subterminal posterior growth zone that contains putative stem cells whose sustained proliferation allows formation of segments during many months [33]. During this phase of juvenile growth, *P. dumerilii* worms also display important regenerative abilities. In particular, after posterior amputation (removal of several segments, the posterior growth zone and the terminal body part bearing the anus named the pygidium), worms are able to regenerate both differentiated structures of the pygidium and stem cells of the growth zone whose activity subsequently allows reformation of the amputated segments [34]. This process, named posterior regeneration, involves formation of a regeneration blastema whose cells likely derive from dedifferentiation of cells belonging to tissues abutting the amputation plane. Indeed, at one and two days post amputation, cells at the amputation site start to express various proliferation and pluripotency stem cell markers [34], suggesting that amputation induces extensive reprogramming of differentiated cells into proliferating progenitor/ stem cells.

Given the well-known importance of epigenetic modifications such as DNA methylation during cellular reprogramming events and evidence for their involvement during vertebrate regeneration *(e.g.,* [3,35–37]), we hypothesized that epigenetic modifications might be important during *P. dumerilii* posterior regeneration and that this process might represent a valuable model to understand how epigenetic regulations influence cellular reprogramming and regeneration. In this study, using *in silico* and experimental approaches, we found high level of CpG methylation in *P. dumerilii* genome, with significant variations during the course of development. Using genomic data and phylogenetic analyses, we identified a full set of *P. dumerilii* writers, modifiers and readers of 5mC methylation, as well as NuRD components. We subsequently studied the evolution of these proteins in animals. We also show that many of the corresponding genes have dynamic expression during development and regeneration. Strikingly, most investigated genes have expression patterns during regeneration similar to those previously documented for stem cell genes [34]. Treatment with a DNA hypomethylating drug, Decitabine (5-aza-2’-deoxycytidine), impairs larval development, regeneration and subsequent segment addition, suggesting a requirement of DNA methylation for posterior regeneration and stem cell-based post-regenerative posterior growth in *P. dumerilii*.

## RESULTS

### High level of CpG methylation in *P. dumerilii*

In a first attempt to characterize DNA methylation in *P. dumerilii*, we used a computational approach that allows evaluating methylation level and pattern of an organism based on determination of normalized CpG content (*e.g.*, [38–40]). Indeed, 5mC is hypermutable through spontaneous deamination, resulting in thymines [41,42]. As 5mC are mostly or exclusively found in CpGs, genomes with high level of 5mC in the germline are therefore predicted to have a low content of CpGs [43]. Determination of CpG observed/expected (o/e) ratio can thus be used as an estimator of 5mC levels, CpG o/e close to 1 means no methylation while CpG o/e far below 1 suggests that methylation of CpGs is present (*e.g.*, [38–40]). As in non-vertebrates methylated CpGs are mostly found within gene bodies [10,11], we calculated CpG o/e for *P. dumerilii* gene bodies, by applying Notos, a software that computes CpG o/e ratios based on kernel density estimations [40,44], on a high quality *P. dumerilii* reference transcriptome [45]. We found a CpG o/e distribution with a single mode at 0.55 (Figure 1A), suggesting high-level gene body methylation in *P. dumerilii.* Indeed, based on a large-scale analysis of 147 species from all major eukaryote lineages, four types of gene body methylation have been defined and *P. dumerilii* fits into type 3 to which belong species with high gene body methylation, for example most vertebrates [44]. We used the same approach to calculate CpG o/e for additional species used for phylogenetic analyses of methylation machinery proteins (Additional file 1: Table S1; Additional file 2: Figure S1) – see below for further discussion.

**Figure 1:**
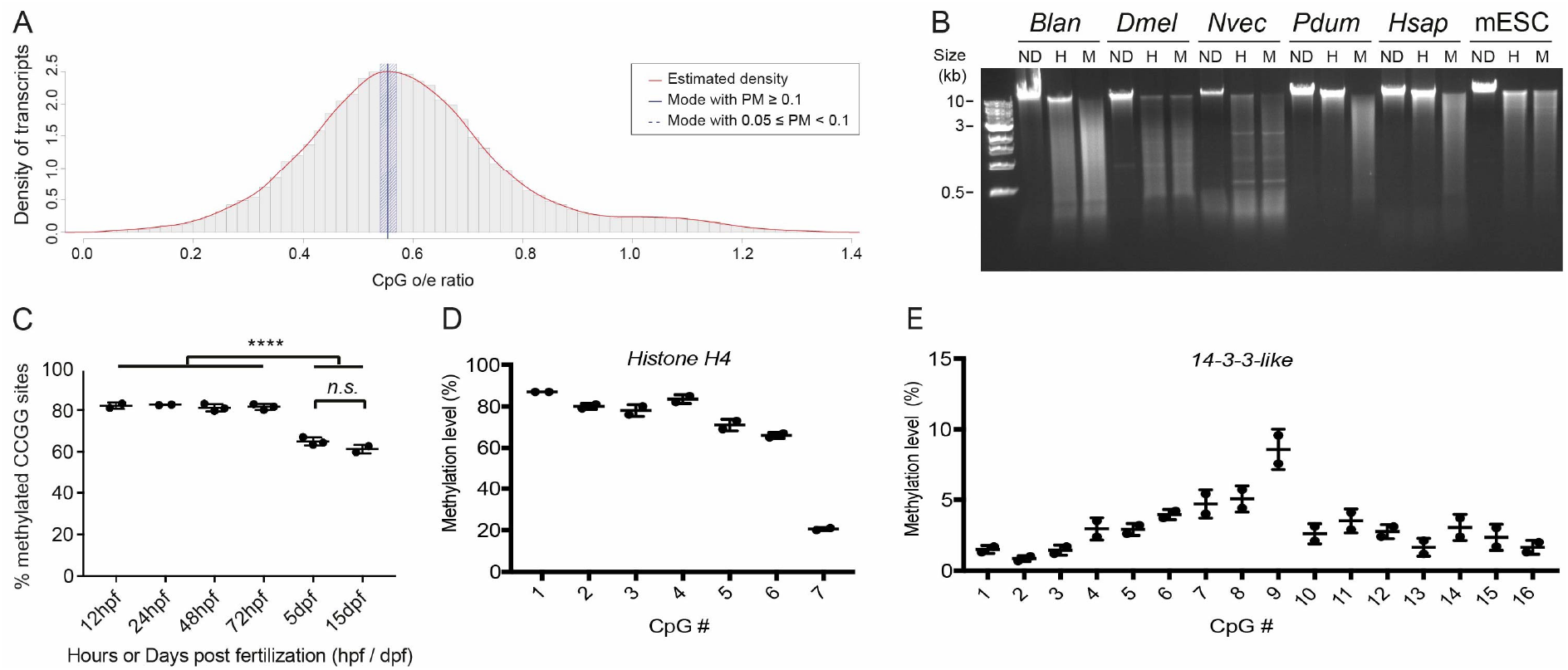
High-level and gene body CpG methylation in *P. dumerilii.* (A) Histogram of CpG o/e ratio of *P. dumerilii* transcripts. Red line indicates the estimated density, vertical blue bar shows estimated mean value and shaded blue bar represents bootstrap confidence intervals of 95%. (B) Electrophoresis of non-digested (ND) genomic DNA (gDNA) or digested with HpaII (H) or MspI (M) from six different animal species with different methylation types. Sizes of fragments, in kilobase pairs (kb), are indicated to the left. Abbreviations: *Blan* = *Branchiostoma lanceolatum; Dmel = Drosophila melanogaster; Nvec* = *Nematostella vectensis; Pdum* = *Platynereis dumerilii; Hsap* = *Homo sapiens;* mESC = *Mus musculus naïve embryonic stem cells.* (C) Graphic representation of DNA methylation measured by LUMA at different stages of *P. dumerilii* life cycle. Mean ± SD. One-way ANOVA, Tukey post hoc test (****: p<0.0001, n.s.: non-significant). (D) and (E) Graphic representation of methylation levels of stretches of CpGs in two *P. dumerilii* genes, *Histone H4* (D) and *14-3-3-like* (E), as defined by bisulfite pyrosequencing. Mean ± SD of two biological replicates is shown.

To further assess CpG methylation in *P. dumerilii* genome at the experimental level, we performed genomic DNA (gDNA) digestion with the methylation-sensitive enzyme HpaII and its methylation-insensitive isoschizomer MspI, which target CCGG sites [46]. If portions of genomes are methylated, different profiles of restriction fragments are expected from the two enzymatic digestions. To facilitate interpretation of profiles obtained with *P. dumerilii* gDNA, we included in our experiment gDNA from species with known methylation patterns (Figure 1B). *Drosophila melanogaster* do not have 5mC methylation and, as previously reported [47], similar profiles are obtained for both HpaII and MspI enzymatic digestions. The cephalochordate *Branchiostoma lanceolatum* and the cnidarian *Nematostella vectensis* have a mosaic pattern of methylation (type 4 in [44]), characterized by the presence of a large number of different cleaved fragments in both digestions and a high-molecular-weight fraction only found with HpaII [46]. Vertebrates such as *Homo sapiens* have global CpG methylation in their genome (type 3 in [44]) and accordingly their gDNA is largely resistant to HpaII digestion [46]. An exception are naïve mouse embryonic stem cells (mESC) [48]; as expected in this case, we found similar restriction profiles with both enzymes. In the case of *P. dumerilii,* we found a restriction pattern that is remarkably similar to that of *H. sapiens,* further supporting high level of CpG methylation in this species (Figure 1B).

We next performed LUminometric Methylation Assay (LUMA) [49,50] to obtain a quantitative assessment of CpG methylation and information about its dynamics during development. LUMA is an efficient method to measure global CpG methylation, based on gDNA digestion (at CCGG sites) by methylation-sensitive restriction enzymes followed by pyrosequencing. LUMA was performed on *P. dumerilii* gDNA extracted from six different developmental stages (Figure 1C). Very high and similar methylation levels were found during embryonic/larval development (from 12 to 72 hours post-fertilization, hpf; about 80% of CCGG sites are methylated). This level significantly decreases after the end of larval development, as shown in juvenile worms (5 and 15 days post-fertilization, dpf), but nevertheless remains quite high (about 60-65% of methylated CCGG sites).

To confirm the existence of gene body methylation in *P. dumerilii,* we performed bisulfite pyrosequencing [51] on CpG-rich parts of the coding region of two different genes, *histone H4* and *14-3-3-like.* These two genes were selected because they display stretches of CpGs in their coding region (7 and 16 CpGs for *histone H4* and *14-3-3-like,* respectively). In addition, orthologs of these genes in other lophotrochozoan species were shown to have gene body methylation [27,29]. Using DNA extracted from 72hpf larvae, we found high levels of methylation (between 65 et 87%) for 6 of the 7 CpGs of *histone H4,* and low levels for all CpGs of *14-3-3-like* (<10%; Figure 1D). These data therefore indicate that gene body methylation does indeed occur in *P. dumerilii* and that level of methylation strongly differs in the two studied genes.

Taken together, these data indicate high-level of CpG methylation in the *P. dumerilii* genome. In addition, 5mC level is dynamic along *P. dumerilii* life cycle and is significantly higher during embryonic/larval development as compared to post-larval stages. We also obtained evidence for gene body methylation in *P. dumerilii* and found that level of CpG methylation in gene bodies is not uniform from one gene to another.

### *P. dumerilii* owns a full ancestral-like DNA methylation and NuRD toolkit

Having established the existence of 5mC in *P. dumerilii,* we next aimed to identify proteins involved in writing, modifying and reading this epigenetic mark, as well as putative NuRD components, in this species. For that purpose, we searched for *P. dumerilii* orthologs of proteins known to exert these functions in mammals, through a sequence-similarity approach, using reciprocal best blasts with *H. sapiens* and *M. musculus* sequences as queries. We found putative *P. dumerilii* orthologs for all investigated proteins/protein families (Additional file 3: Figure S2). Sequences of all the identified proteins can be found in Additional File 4. As 5mC and NuRD proteins are often characterized by the presence of particular domains or association of domains, we searched for conserved domains present in the retrieved *P. dumerilii* proteins. We found in most cases domains that are consistent with orthology relationships inferred from blast analyses (Additional file 3: Figure S2).

As defining orthology relationships only on blast analyses can be misleading, in particular in the numerous cases in which there are several paralogs, we turned out to phylogenetic analyses to ascertain these relationships. To do these analyses on a firm basis and to get insight into the evolution of the DNA methylation and NuRD toolkit in animals, we retrieved, by reciprocal blast searches using mouse and human sequences as queries, putative orthologs from 51 additional species from diverse phylogenetic groups. We ended up with a sample of 54 species from all major animal lineages (Figure 2). Maximum-Likelihood (ML) trees were constructed for each protein family and are shown in Additional file 5: Figure S3. These phylogenetic trees allow to confirm orthology relationships for all *P. dumerilii* proteins and to define the number of members of all protein families in the 54 investigated species (Figure 2). We also summarized the number of members for each defined family. Based on parsimony, we also inferred the presence or absence of each protein family at key nodes of the animal phylogenetic tree and mapped the main gene duplication and gene loss events (Figure 3). All the identified proteins are listed in Additional file 1: Table S2 and their sequence can be found in Additional file 4.

**Figure 2:**
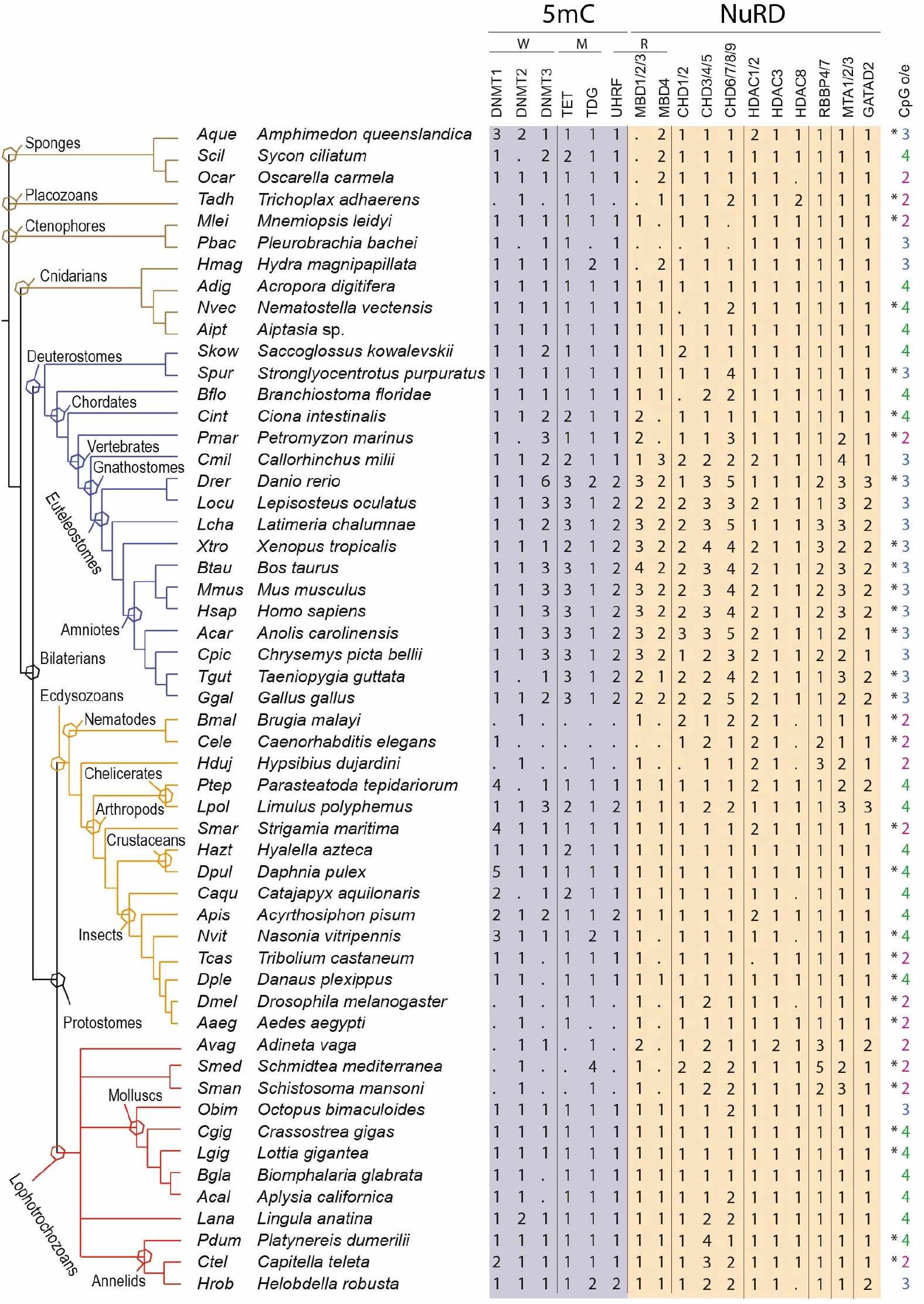
DNA methylation and NuRD toolkit in metazoans. On the left is shown a phylogenetic tree of the 54 metazoan species for which we identified DNA methylation and NuRD genes. Branches of this phylogenetic tree are color-coded (brown for non-bilaterians, blue for deuterostomes, orange for ecdysozoans and red for lophotrochozoans). Polytomies highlight uncertainties about the relationships between bilaterian and non-bilaterian groups and within lophotrochozoans. Major phylogenetic groups are shown by hexagons placed on tree nodes defining these groups. Additional taxonomic information: *Saccoglossus kowaleskii* belongs to hemichordates, *Stronglylocentrotus purpuratus* to echinoderms, *Branchiostoma floridae* to cephalochordates, *Ciona intestinalis* to urochordates, *Hypsibius dujardini* to tardigrades, *Adineta vaga* to rotifers, *Schmidtea mediterranea* and *Schistosoma mansoni* to platyhelminthes, and *Lingula anatina* to brachiopods. Number of members found for each gene families are indicated for each species and dots indicate that we failed to identify any members. Chd1/2, Chd6/7/8/9 and Mbd4 gene families which do not encode NuRD members, are also indicated. CpG o/e clusters [44] are also shown (stars indicate data derived from a previous study [44]). W = Writers, M= Modifiers, R = Readers.

**Figure 3:**
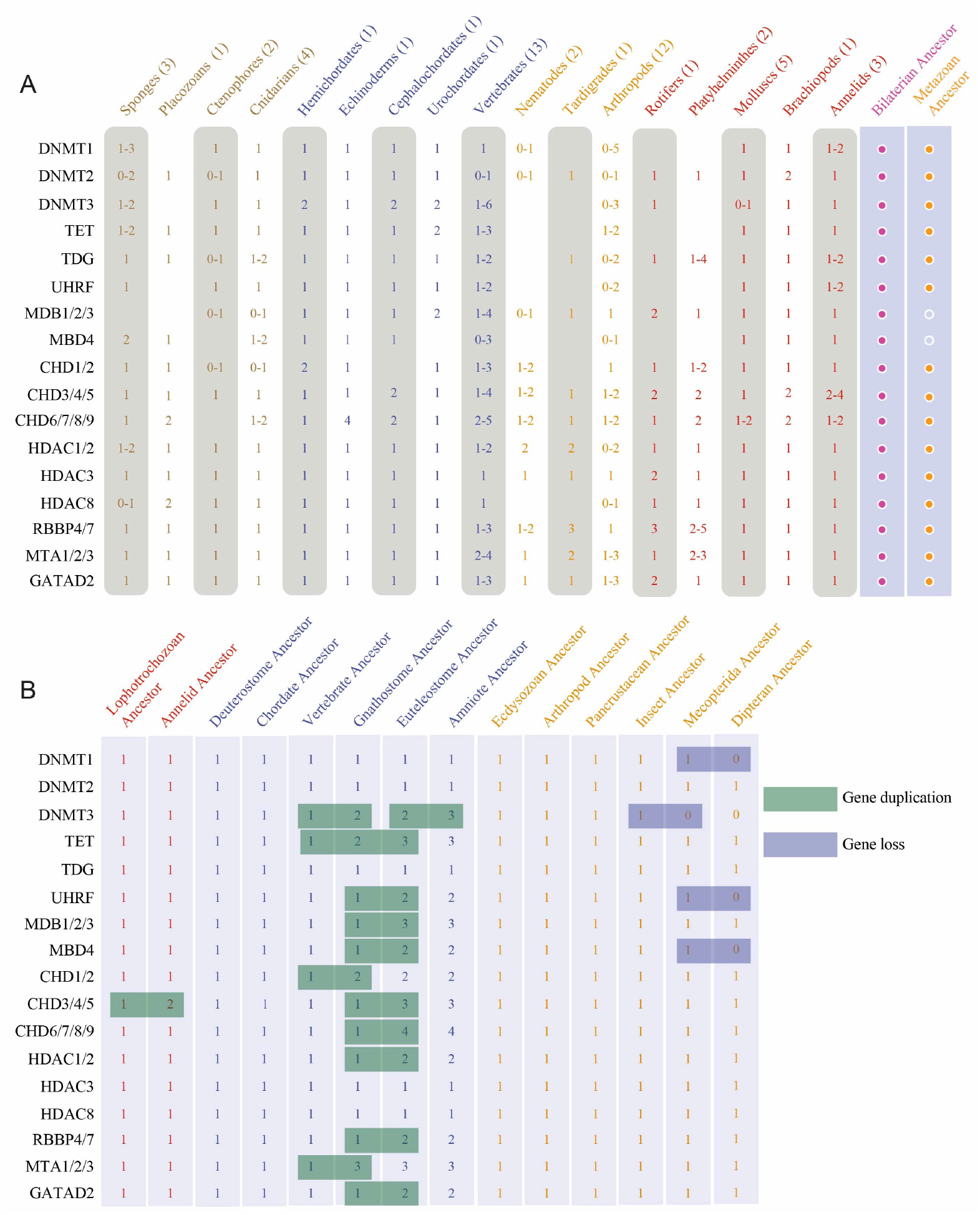
Evolution of DNA methylation and NuRD gene families in metazoans. (A) The number (or range of numbers) of members of each family/subfamily in the indicated phylogenetic groups is shown (none if no member detected). The number of studied species in each phylogenetic group is indicated next to the group name. Two final columns summarize the putative ancestral set of all studied families/subfamilies in metazoan and bilaterian ancestor. Putative ancestral set of families/subfamilies in eumetazoan ancestor is the same than the one for bilaterian ancestor and is not shown for sake of clarity. (B) Putative numbers of members of all studied families/subfamilies in the last common ancestor of selected phylogenetic groups are shown. These numbers have been inferred using both the number of members in studied species of these groups and phylogenetic analyses. Green boxes highlight increases in number of members, thus gene duplications; blue boxes highlight decreases in number of members, thus gene loss events. Most represented phylogenetic groups are those indicated on the phylogenetic tree of Figure 2 or listed in its legend, except pancrustaceans (crustaceans + insects), Mecopterida (which includes dipterans and lepidopterans) and dipterans (which include *D. melanogaster* and *A. aegypti).*

*P. dumerilii* genome encodes three Dnmt proteins that can be clearly assigned to the Dnmt 1, 2 and 3 subclasses (Additional file 5: Figure S3A). The presence of these three subfamilies appears to be ancestral to animals (Figure 3), as these three subfamilies are found in most non-bilaterians and in many species in the three bilaterian evolutionary lineages (Figure 2). Only few gene duplications occurred for *dnmt1* (in particular in some arthropod species) and for *dnmt3* (in particular two duplications during vertebrate evolution). *dnmt*gene losses occurred in some species or lineages such as nematodes, rotifers, tardigrades, and Platyhelminthes (Figure 3). Absence of both *dnmt1* and *dnmt3* is correlated to the absence or very low abundance of cytosine DNA methylation as shown by CpG o/e ratio calculation (Figure 2). *P. dumerilii* also owns single *tet, tdg* and *uhrf* genes, which likely corresponds to the ancestral situation in animals (Figures 2 and 3; Additional file 5: Figure S3B-D).

Duplications of *tet* genes are unfrequent, but two duplications nevertheless occurred in vertebrates, likely one before diversification of gnathostomes and the second before that of euteleostomes (bony vertebrates; Figure 3). *tet* genes are only absent in species that lack 5mC methylation and *dnmt1* and 3, with the exception of dipterans which possess one *tet* gene. *tdg* is present in almost all investigated species in one copy, as expected for a gene involved in DNA repair. *uhrf* has a similar distribution that *tet,* being absent in species lacking 5mC, but in this case including dipterans. A single *uhrf* gene is found in most other species, with the notable exception of euteleostomes that possess two genes, indicating the occurrence of a duplication in this evolutionary lineage (Figure 3).

Phylogenetic analysis shows the existence of two large groups of Mbd proteins, one which contains Mbd1, Mbd2 and Mbd3 proteins from vertebrates (hereafter named Mbd1/2/3 group) and the other which contains vertebrate Mbd4 and MeCP2 proteins (Mbd4 group; Additional file 5: Figure S3E). *P. dumerilii* genome encodes two Mbd proteins, one belonging to Mbd1/2/3 group (putative NuRD component) and the other to Mbd4 group. Presence of both Mbd1/2/3 and Mbd4 is observed in most species in the three bilaterian lineages, as well as in most cnidarians, indicating that the last common ancestor of bilaterians and cnidarians (eumetazoans) owned at least two *mbd* genes (Figures 2 and 3). It is not clear, in contrast, whether the last common ancestor of all animals possessed one or two *mbd* genes, as Porifera and Ctenophora, which have been both proposed as the potential sister group of all other metazoans, have members only of Mbd4 or Mbd1/2/3 class, respectively (Figure 3). The presence of a single *mbd* gene may therefore represent the ancestral situation in metazoans. *mbd* gene losses occurred in few species, mainly in those that also lack cytosine DNA methylation. Few gene duplications also occurred, in particular in vertebrates in which both *mbd1/2/3* and *mbd4* ancestral genes underwent gene duplications (Figure 3).

Phylogenetic tree of Chd proteins comprises three large groups, one that includes vertebrate Chd3/4/5 proteins (hereafter named Chd3/4/5 group), the second vertebrate Chd1/2 (Chd1/2 group), and the third vertebrate Chd6/7/8/9 (Chd6/7/8/9 group; Additional file 5: Figure S3F). Six *chd* genes have been found in *P. dumerilii,* one belonging to the Chd1/2 group, one to Chd6/7/8/9 group and four to the Chd3/4/5 group (putative NuRD components). Member of these three groups are found in almost all studied species, including non-bilaterians, indicating that presence of three different types of CHD proteins is ancestral to animals (Figures 2 and 3). Only very few gene losses occurred. Gene duplications are more frequent, in particular in vertebrates and lophotrochozoans, including in annelids in which two to four members are found in the three studied species (Figure 3).

Previous phylogenetic studies classified Hdac proteins into four classes (I, IIA/B, III and IV) [52,53]. Here we focused on class I to which belong *Hdac1, Hdac2, Hdac3,* and *Hdac8* genes, which encode members of the NuRD complex. Phylogenetic analysis showed the existence of three subgroups, Hdac1/2, Hdac3, and Hdac8 (Additional file 5: Figure S3G). We found one member of each subgroup in *P. dumerilii,* as well as in almost all other investigated species, indicating that at least three class I *hdac* genes were present in the last common ancestor of all animals (Figures 2 and 3). We found only very few gene losses *(e.g.,* in nematodes). Duplications mainly occurred in arthropods and vertebrates. Single *rbbp4/7, mta1/2/3* and *gatad2* genes are found in *P. dumerilii* (Additional file 5: Figure S3 H-J). At least one member of each of these subfamilies is found in all studied species, indicating that their presence is ancestral to animals (Figures 2 and 3). Gene duplications occurred in vertebrates, ecdysozoans and lophotrochozoans (Figure 3).

In conclusion, we identified a complete set of writers, modifiers and readers involved in 5mC methylation, as well as putative NuRD components, in *P. dumerilii.* In addition, we provide an animal-wide view of the evolution of the corresponding gene families (Figure 2), which suggests that the last common ancestor of animals already owned a complex repertoire of 5mC and NuRD toolkit genes (Figure 3). Our analysis also indicates that *P. dumerilii* repertoire is mostly composed of single-copy genes and likely close to the one present in the last common ancestor of bilaterians.

### DNA methylation and NuRD tookit genes are dynamically expressed during development and regeneration in *P. dumerilii*

We next aimed to characterize expression of DNA methylation and NuRD genes in *P. dumerilii*. We first took advantage of two previously published transcriptomic datasets corresponding to various developmental stages and adult condition of *P. dumerilii* [45,54], available in a public database (Pdumbase) [55]. The first dataset corresponds to embryonic developmental stages, ranging from 2 to 14hhpf, with a time point every two hours [45]. The second dataset comprises major larval stages (24hpf to 4dpf; five time points), juvenile stages (10dpf to 3 months post fertilization (mpf); five time points) and adult reproductive stages (males and females) [54]. Altogether expression data for a total of 19 stages during the course of embryonic and post-embryonic development, as well as male and female adult stages, are available.

Expression values for most genes (exceptions are *Pdum-dnmt3* not found in the two sets of transcriptomic data and *Pdum-gatad2* and *Pdum-rbbp4/7* only found as chimeric transcripts) were recovered and can be found in Additional file 6: Figure S4. High transcript levels are found for many studied genes in the earliest developmental stages (2-6hpf) and several of them belong to co-expression clusters defined by Chou et al. [45] as maternal gene clusters (clusters 1-4; Additional file 6: Figure S4). This indicates that the *P. dumerilii* egg contains a large pool of maternal transcripts coding for DNA methylation proteins that could be used for embryonic development. To further analyze these expression data, we studied changes in expression during main steps of *P. dumerilii* life cycle (Figure 4).

**Figure 4:**
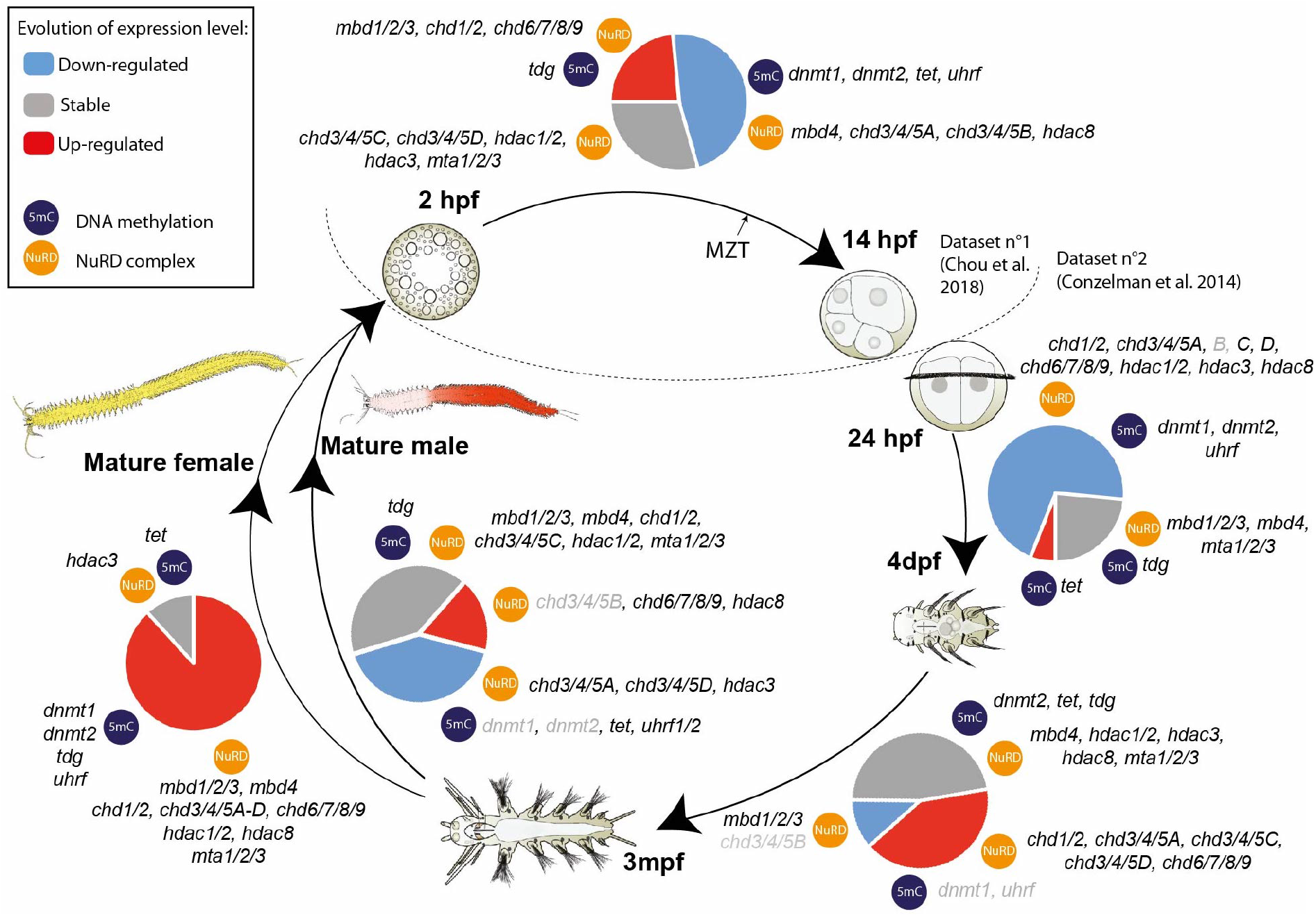
DNA methylation and NuRD genes have dynamic expressions during *P. dumerilii* life cycle. Main steps of *P. dumerilii* life cycle are shown. Expression changes between consecutive stages are indicated as “Down-regulated” (expression level decreases with a fold change superior to two), “Stable” (fold change inferior to 2) and “Up-regulated” (expression level increases with a fold change superior to two). Proportions of these three categories are represented as color pie charts. Genes belonging to 5mC machinery and NuRD/NuRD-related are mentioned. Grey letterings indicate genes with low expression (< 5 Fragments Per Kilobase Million, FPKM) in the compared stages. The two datasets that have been used are indicated [45,54]. Maternal to zygotic transition (MZT) is shown at around 10hpf as previously suggested [45]. hpf = hours post-fertilization, dpf = days postfertilization, mpf= months post-fertilization. For sake of clarity, *“Pdum-”* prefixes have been omitted. Drawings of embryos, larvae and worms are adapted from [32].

From 2hpf to 14hpf, a decrease in quantity of transcripts of about half of the genes, including genes coding for DNA methylation maintenance *(Pdum-dnmt1* and *Pdum-uhrf),* as well as putative members of NuRD complex *(Pdum-chd3/4/5A-B* and *Pdum-hdac8),* is observed. From 24hpf to 4dpf, this decrease is found for most genes, including *Pdum-dnmt1* and *Pdum-uhrf*. In contrast, expression of *Pdum-tet* and *Pdum-tdg* is increased or stable, respectively. This is consistent with the decrease of level of CCGG methylation that we observed at the end of larval development (Figure 1C). From 4dpf to 3mpf, a majority of genes have stable expression with the exception of all but one *P. dumerilii chd* genes that are up-regulated (Figure 4). Transition from 3mpf to adult stage is strikingly gender-specific: in males most genes have stable or down-regulated expressions, while in females about 80% of the genes are strongly up-regulated (Figure 4; Additional file 6: Figure S4), suggesting different occurrence and importance of DNA methylation during sexual maturation and gamete production between males and females.

We next studied expression of DNA methylation and NuRD genes during *P. dumerilii* posterior regeneration. Using RNA-seq data (Gazave et al., unpublished), we found most of these genes to be expressed throughout regeneration (Additional file 1: Table S3). No clear trend in the evolution of their expression can however be detected from these data. To gain more insights into the expression during regeneration of DNA methylation and NuRD genes and characterize in which part(s) and tissue(s) of the regenerated region these genes are expressed, we performed whole-mount RNA *in situ* hybridizations (WMISH) at different stages of posterior regeneration [34], focusing on a set of ten genes that encode putative writer/modifier/readers of 5mC or NuRD components, and/or show dynamic expression in RNA-seq data. Representative expressions are shown in Figure 5 and Additional file 7: Figure S5. Schematic representation of the expression patterns can be found in Additional file 8: Figure S6. *Pdum-dnmt1* and *Pdum-dnmt3* are weakly expressed in the wound epithelium at stage 1 (Figure 5 A, B). At stage 2, *Pdum-dnmt1* is strongly expressed in two internal groups of cells and in the lateral ectoderm (Figure 5 A’). Its expression extends almost in the whole blastema at stage 3 and is found in the regenerated growth zone (Figure 5 A’’). At the same stages, *Pdum-dnmt3* is very weakly expressed in both mesodermal and ectodermal cells of the regenerated region (Figure 5 B’, B’’). At stage 4 and 5, *Pdum-dnmt1* is expressed in mesoderm of the developing segments, mesodermal and ectodermal growth zone, at the basis of the anal cirri and weakly in the lateral/dorsal ectoderm (Figure 5 A’’’, A’’’’; Additional file 7: Figure S5A). *Pdum-dnmt3* is expressed in the ventral ectoderm and at the basis of the anal cirri (Figure 5 B’’’, B’’’’). *Pdum-tet* expression is not trustfully detected at stage 1 (Figure 5C). At stage 2 and 3, weak expression is found in both internal and superficial blastemal cells (Figure 5C’-C’’), which continues at stage 4 and 5 at which expression at the basis of anal cirri is also observed (Figure 5 C’’’, C’’’’). *Pdum-tdg* is strongly expressed at stage 1 in the wound epithelium and internal cells of the segment abutting the amputation plane (Figure 5D). At stage 2 and 3, it is broadly expressed in the whole blastema (Figure 5D’, D’’). From stage 3, *Pdum-tdg* is expressed in mesoderm and ectoderm of the developing segments, and in ectodermal and mesodermal growth zones, as well as weakly at the basis of the anal cirri (Figure 5 D’’’, D’’’’; Additional file 7: Figure S5B).

**Figure 5:**
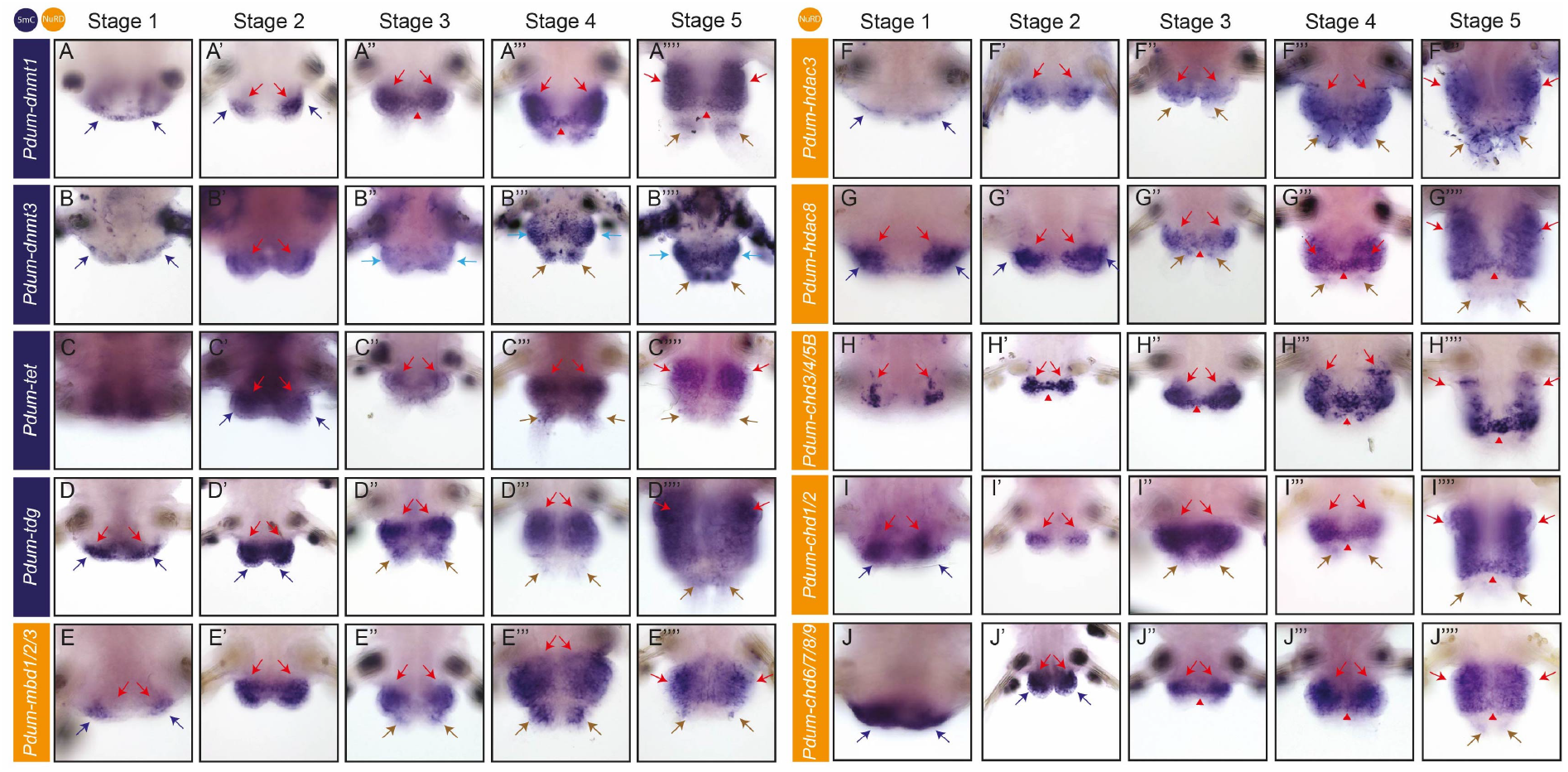
DNA methylation and NuRD genes are expressed during most or all stages of *P. dumerilii* regeneration. Expression patterns obtained by whole-mount *in situ* hybridization (WMISH) for genes whose name is indicated on the left at the five previously defined stages of posterior regeneration [34] are shown. All panels are ventral views (anterior is up). In stage 1, dark blue arrows point to the wound epithelium and red arrows to internal cells of the segment adjacent to the amputation. In the other stages, dark blue arrows point to epithelium covering the blastema, red arrows point to mesodermal cells of the blastema or developing segments (depending on stages), red arrowheads to the mesodermal part of the growth zone, light blue arrows to ectodermal expression, including segmental ectodermal stripes, and brown arrows to the basis of anal cirri.

*Pdum-mbd1/2/3, Pdum-hdac3* and *Pdum-hdac8* have roughly similar expression during posterior regeneration, *Pdum-hdac3* being expressed at most stages weaker than the two other genes. At stage 1, an expression is detected in two lateral patches of cells in and close to the wound epithelium (Figure 5E, F, G). The three genes are widely expressed in the blastema at stage 2 and 3 (Figure 5E’, E’’, F’, F’’, G’, G’’). Expression in mesoderm and ectoderm of the developing segments, growth zone and at the basis of the anal cirri is observed at stage 4 and 5 (Figure 5 E’’’, E’’’’, F’’’, F’’’’, G’’’, G’’’’; Additional file 7: Figure S5C, D). *Pdum-chd3/4/5B* expression is found at stage 1 in four small patches of internal cells close (but not adjacent) to the wound epithelium, two located ventrally and two dorsally (Figure 5H; Additional file 7: Figure 5E). From 2dpa, we observed an intense expression in the mesodermal part of the regenerated region, including the mesodermal growth zone (Figure 5H’-H’’’’). *Pdum-chd1/2* and *chd6/7/8/9* are expressed in cells in and close to the wound epithelium at stage 1, the latter having a much broader expression (Figure 5I, J). At stage 2, both genes are expressed in superficial and internal cells of the regenerated region, in most or all cells for *Pdum-chd6/7/8/9* but only in a few cells for *Pdum-chd1/2* (Figure 5I’, J’). Broad expression in mesodermal cells, including growth zone, is observed at later stages (Figure 5I’’, I’’’, I’’’’, J’’, J’’’, J’’’’). At stage 5, *Pdum-chd1/2* is also weakly expressed in the ectodermal growth zone (Additional file 7: Figure 5F).

Altogether, we found that that *P. dumerilii* DNA methylation and NuRD genes are dynamically expressed during embryonic, larval and post-larval development, as well as during sexual maturation and regeneration. During this latter process, most genes are expressed from its earliest stages and their expression is later mostly found in blastemal cells, putative mesodermal and ectodermal stem cells of the growth zone and cells of the developing segments (Additional file 7: Figure 6). Observed patterns of expression show striking similarities with those previously reported for proliferation and stem cell genes [34], suggesting that DNA methylation and NuRD genes are mainly expressed in undifferentiated proliferating cells, including stem cells of the regenerated posterior growth zone.

**Figure 6:**
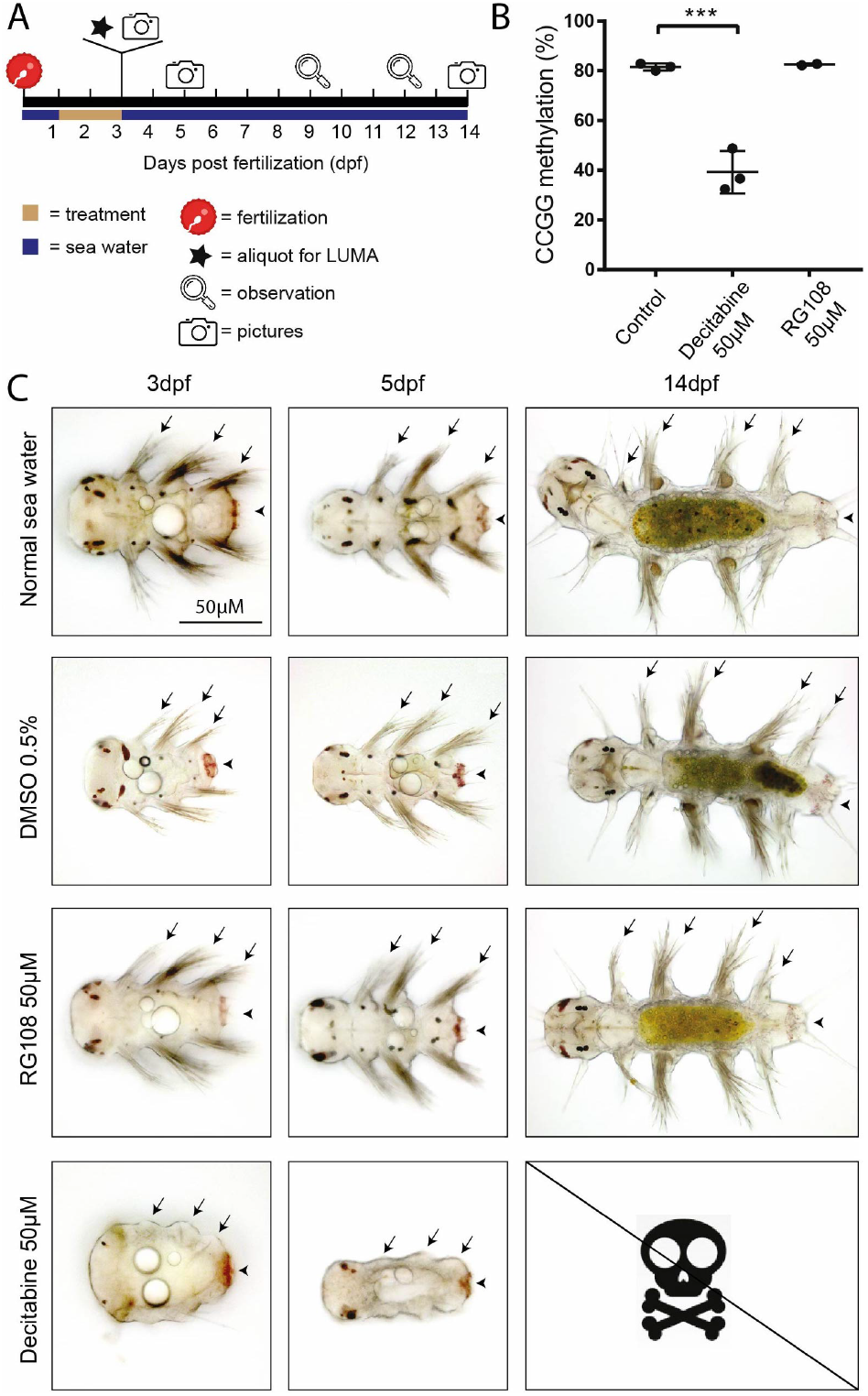
Decitabine treatment decreases DNA methylation level and impairs larval development. (A) Schematic representation of the experimental design. Larvae were treated with Decitabine (50μM), RG108 (50μM) or DMSO (0,5 %; control) from 1 day post-fertilization (dpf) to 3dpf. At 3dpf, a part of the larvae was frozen for subsequent DNA methylation measurement with LUMA and remaining larvae were placed and kept in normal sea water until 14dpf. Observations were done at indicated time points and pictures taken at 3, 5 and 14dpf. (B) Graphic representation of CCGG DNA methylation as measured by LUMA for the different conditions. Mean ± SD. One-way ANOVA, Dunnet post hoc test was performed (***: p < 0.001). (C) Morphological observations at 3, 5 and 14dpf. Ventral views of representative larvae/juvenile worms are shown (anterior on the left). At the three time points, RG108-treated larvae/juvenile worms show morphology similar to that of controls (sea water and DMSO 0.5 %). At 14dpf, like control worms, RG108-treated have added a fourth segment. In contrast, Decitabine-treated larvae/juvenile worms display at 3 and 5dpf an abnormal morphology with strongly reduced appendages (parapodia; arrows) and reduced pygidium (arrowheads). Massive death occurred in the 5 to 9dpf time period, so no Decitabine-treated worms could be observed at 14dpf.

### Decitabine reduces DNA methylation and impairs development, regeneration and post-regenerative posterior growth in *P. dumerilii*

To test a possible role of DNA methylation during *P. dumerilii* regeneration, we tried to reduce 5mC level using two well-known and widely-used hypomethylating agents, Decitabine (5-aza-2’-deoxycytidine) and RG108 (N-Phthalyl-L-Tryptophan) [56–58]. Decitabine is incorporated in DNA and binds Dnmt1 irreversibly, leading to progressive loss of DNA methylation through cell divisions. RG108 is a specific non-nucleoside inhibitor of Dnmt1, which acts by binding in a reversible manner to the active center of the enzyme. As these two drugs have never been used in *P. dumerilii,* we first tested their activity by treating larvae continuously from 1 to 3dpf with Decitabine or RG108 (Figure 6A).

Neither drugs caused significant lethality during treatment. DNA was extracted from larvae at 3dpf and CCGG methylation level measured using LUMA (Figure 6B): Decitabine treatment leads to a 2.5-fold decrease of CCGG methylation (from 81.5% to 32.4%) while no significant effects were found for RG108. We also checked for morphological defects (Figure 6C): larvae were observed either immediately after treatment (at 3dpf) or after the drug having been washed out and larvae put in normal sea water until 5 or 14dpf. Larvae that have been treated with Decitabine present morphological abnormalities at 3dpf, in particular reduced parapodia (worm appendages) bearing very few chaetae (extracellular chitinous structures) and a reduced pygidium (Figure 6C). While abnormal, these larvae were alive and survived for a few more days. All animals did however die in the following days, possibly because of feeding defect (during this period normal young worms start to eat, dead Decitabine-treated worms consistently showed an empty gut). In contrast, RG108 treatment did not affect larval morphology (Figure 6C). We therefore conclude that Decitabine can affect DNA methylation level and larval development in *P. dumerilii.*

To investigate potential consequences of a decrease of DNA methylation on regeneration, we treated worms with three concentrations of Decitabine (10μM, 50μM and 100μM) immediately after amputation for five days and scored the worms every day for the stage that has been reached based on a previously established staging system [34]. We found a small number of deaths at 10μM and 50μM concentrations, while 100μM concentration appears to be much more harmful to worms (Additional file 1: Table S4). Some worms also underwent spontaneous amputation of their posterior part (autotomy) at some time points (Additional file 1: Table S4) and these worms were excluded from the analysis. We found that Decitabine significantly delayed regeneration as compared to controls (DMSO 0.5 % and sea water), in a concentration-dependent manner (Figure 7A). At 5 days post-amputation (dpa), while the vast majority of control worms reached stage 4 or more, worms treated with Decitabine were mostly at stages 2 to 3 (Figure 7B). No major abnormalities were observed at the morphological level in Decitabine-treated worms (not shown). To better understand how regeneration proceeds in presence of Decitabine, we did Decitabine treatment (at 50μM as this concentration shows low toxicity and important effect on regeneration) from 0dpa to 5dpa, fixed treated worms at 5dpa and performed WMISH for some of the genes whose expression was previously studied during normal regeneration [34]. The analyzed genes showed expressions at 5dpa in Decitabine-treated worms which are similar to those of stage 2 or 3 in non-treated worms [34], indicating that regeneration is blocked in presence of Decitabine (Figure 7C). Abnormal expressions, never observed in non-treated animals, were nevertheless found in some treated worms for *Pdum-hox3* (growth zone marker; extended and/or mis-located expression domain), *Pdum-piwiB* (stem cell marker; no or reduced expression), and *Pdum-engrailed* (segment marker; incomplete expression stripes).

**Figure 7:**
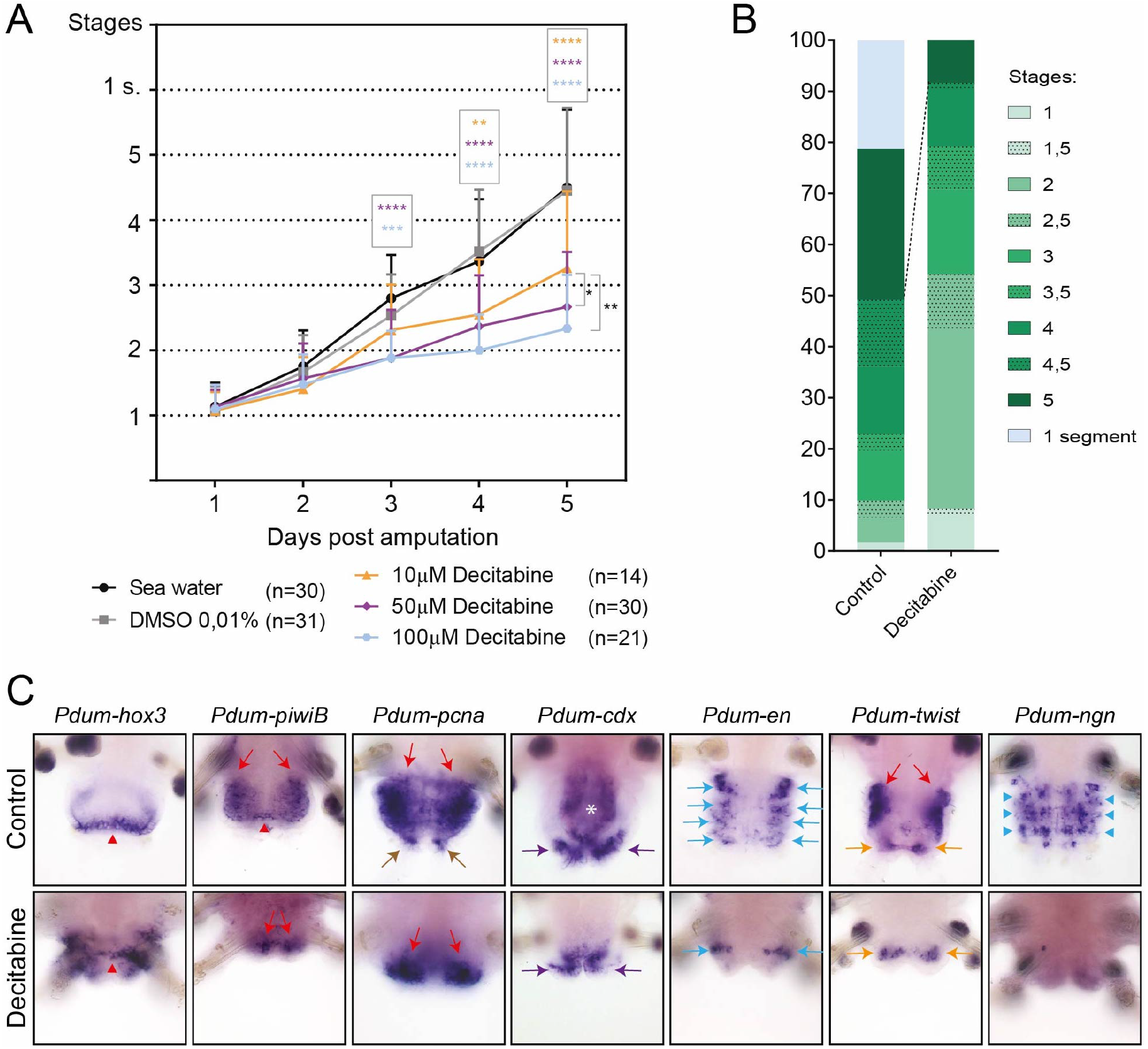
Decitabine treatment impairs posterior regeneration. (A) Graphic representation of the stages reached by control worms (normal sea water and DMSO 0,01%) and worms treated with three different concentrations of Decitabine every day for five days. Regeneration is delayed for 50μM and 100μM Decitabine conditions, from 3 days post-amputation (dpa) onwards, and for 10μM Decitabine conditions, from 4dpa onwards. Three experiments, mean ± SD. 2-way ANOVA (p-value: Time p < 0.0001, Treatment p < 0.0001, Interaction p < 0.0001) with Tukey post hoc test (**: p<0.01; ***: p < 0.001; ****: p< 0.0001). (B) Proportions of control and Decitabine-treated worms with different regeneration scores at 5dpa. Worms treated with the three different concentrations of Decitabine have been pooled. Most worms treated with Decitabine did not reach stage 5 at 5dpa, while about 50% of control worms reached this stage (some of which already having produced a first new segment). (C) Ventral views of WMISH at 5dpa of posterior part of control worms and worms treated with 50μM Decitabine are shown. Expression of marker genes for different structures/tissues/cells during posterior regeneration has been studied, *i.e., Pdum-hox3* (growth zone), *Pdum-piwiB* (growth zone and segmental progenitors), *Pdum-pcna* (proliferating cells), *Pdum-caudal/cdx* (pygidium and growth zone), *Pdum-engrailed (Pdum-en;* segmental stripes), *Pdum-twist* (muscle progenitors), and *Pdum-neurogenin (Pdum-ngn;* neural progenitors) [34]. Red arrowheads point to expression in the growth zone, red arrows to mesodermal expression, brown arrows to expression at the basis of the anal cirri, violet arrows to expression in pygidium, light blue arrows to segmental ectodermal expression, light blue arrowheads to neural progenitors, and orange arrows to expression in pygidial muscles. White asterisks indicate expression in the hindgut.

It has been shown that mammalian cells treated with Decitabine only partially recover their initial methylation level, leading to an epigenetic ‘imprint’ of drug exposure [59]. We hypothesized that Decitabine treatment could have long-term impacts in *P. dumerilii* and affect segment formation that follows regeneration (post-regenerative posterior growth [33,34]).

To test this hypothesis, we treated worms with Decitabine from 0 to 5dpa, then washed out the drug, put worms in normal sea water until 25dpa, checking their morphology and counting the number of segments that have been produced at six time points (Figure 8A). As for the previous experiment, Decitabine treatments induced few worm deaths and autotomies (Additional file 1: Table S5). Most Decitabine-worms recovered from the treatment and were able to reach stage 5 and undergo posterior growth (Figure 8B). Decitabine-treated worms continued to be delayed as compared to controls worms, had a reduced number of newly added segments at 25dpa (treated worms owned about 4 to 6 segments compared to about 10 to 12 segments for controls), and showed morphological abnormalities (Figure 8B, C). Reduced number of newly added segments was due not only to a marked delay during regeneration, but also to a reduced rate of segment addition after the drug had been washed out (Additional file 9: Figure S7A). A high variability was observed among Decitabine-treated animals compared to controls (Additional file 9: Figure S7B-F). We defined three classes of animals based on their morphology and the number of newly added segments at 25dpa (Figure 8C; Additional file 10: Figure S8). Class 1 animals (30.6% of Decitabine-treated worms) show a characteristic bottleneck-like shape with a marked constriction between non-regenerated and regenerated regions, no or few newly added segments, and no or abnormal anal cirri. These worms are prone to undergo autotomy. Class 2 animals (61.6%) have an abnormal shrink-like shape, a reduced number of newly added segments, absence of well-differentiated parapodia on newly added segments, and no or abnormal anal cirri. Class 3 worms (7.9%) have a morphology and number of newly added segments similar to non-treated animals.

**Figure 8:**
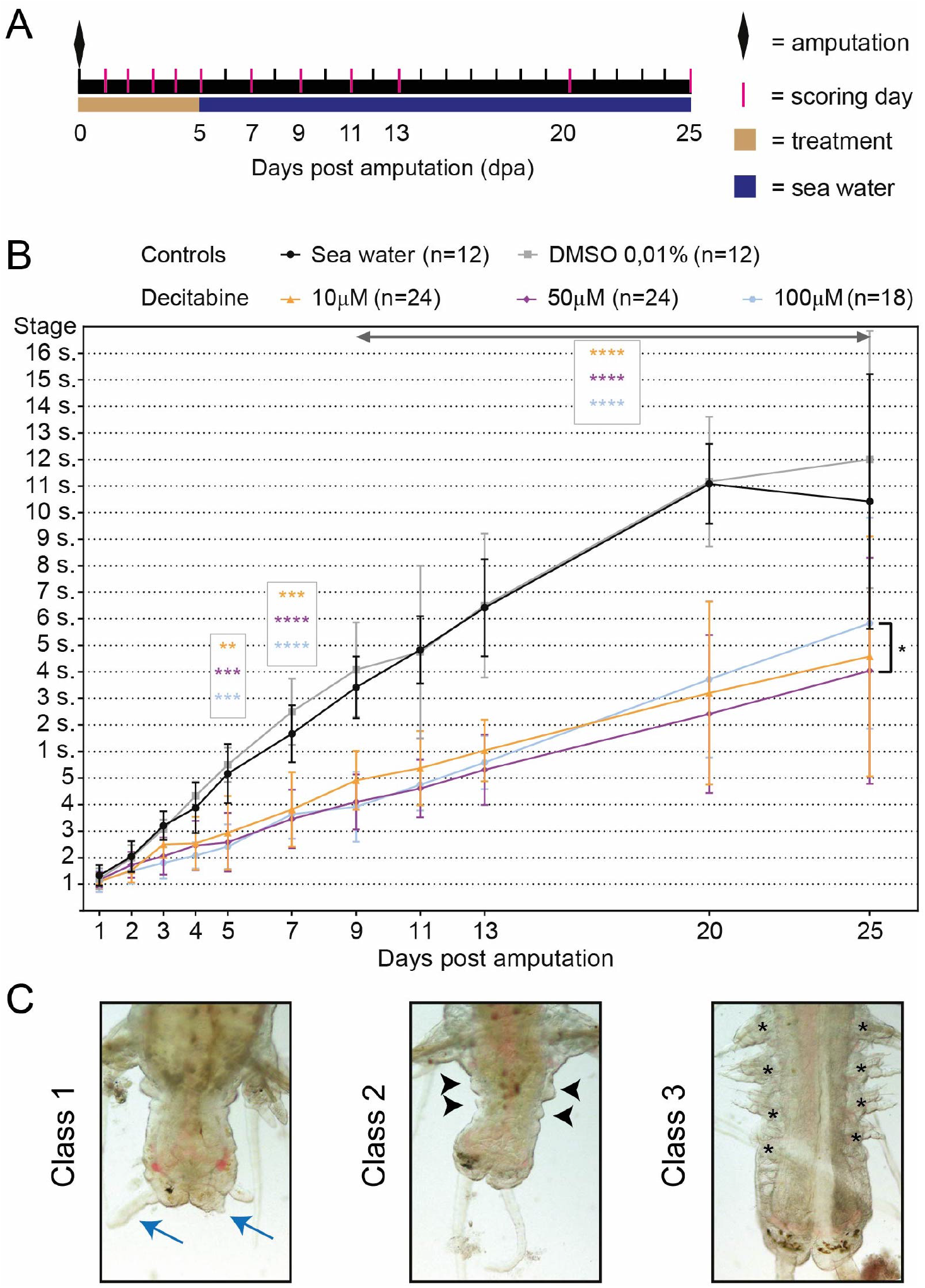
Decitabine treatment during posterior regeneration affects subsequent post-regenerative posterior growth. (A) Schematic representation of the experimental design. Worms were treated with Decitabine (10μM, 50μM or 100μM) or from amputation to 5 dpa. Control animals are treated with DMSO 0,01% or put in normal sea water. After washing out, worms were kept from 5dpa to 25dpa in normal sea water and observed at several time points until 25dpa. (B) Graphic representation of regeneration stage that has been reached or number of newly added segments by Decitabine-treated and control worms. A significant delay in post-regenerative posterior growth is observed in Decitabine-treated worms as compared to controls. Two experiments, mean ± SD, 2-way ANOVA (p-value: Time p < 0.0001, Treatment p < 0.0001, Interaction p < 0.0001) with Tukey post hoc test (**: p<0.01; ***: p < 0.001; ****: p< 0.0001). Only p-values corresponding to comparison to normal sea water are shown, as highly similar ones are obtained for comparison to DMSO controls. (C) Representative morphology at 25dpa of worms belonging to the three defined classes (see main text for details). While class 3 worms showed well-differentiated segments with parapodia (black asterisks), no or reduced parapodia were observed in class 1 and class 2 (black arrowheads) worms, respectively. Small or abnormally shaped anal cirri (blue arrows) were frequently observed in class 1 and 2 worms.

Taken together, these observations indicate that Decitabine treatment during regeneration has long-term effects and affects subsequent post-regenerative posterior growth, possibly by affecting growth zone regeneration. Some Decitabine-treated worms were however able to add new segments in an almost normal manner, which led us to hypothesize that the growth zone was not impacted in a similar manner in all animals. To point out a potential link between regeneration of the growth zone and ability to later add segments, we performed a multiple correlation analysis (Additional file 11: Figure S9). In control worms, as expected, only positive correlations were observed, which means that, for example, worms with numerous segments at 20dpa already had a high number of segments at 11dpa. In contrast, in Decitabine-treated worms, while there were positive correlations for closely related days of scoring (for example: 2 to 3dpa, 3 to 4dpa, …), negative correlations were also found and suggested that treated worms that regenerated faster eventually produced less segments. It has been shown that the growth zone is regenerated and become functional, producing news segments, at about 3dpa [34]. Our interpretation is therefore that worms with high regeneration scores (scored at stage 3 or more) at 5dpa regenerated a dysfunctional growth zone in the presence of Decitabine, which later led to a null or reduced production of segments, the few produced ones having in addition morphological abnormalities. Worms with low scores (less than 3) probably did not regenerate their growth zone during Decitabine treatment period and did it after 5dpa in the absence of the drug, which led to the formation of a functional growth zone and therefore to normal segment addition. Our data therefore suggest that Decitabine affects the functionality of the growth zone. Consistently, expression of growth zone, stem cell and segment markers (*Pdum-hox3*, *Pdum-piwiB* and *Pdum-engrailed,* respectively) is affected in some Decitabine-treated worms (Figure 7C).

As described above, worms treated with Decitabine from 0 to 5dpa show morphological abnormalities at 25dpa. To better understand these alterations, we performed WMISH on Decitabine-treated worms at 25dpa for a set of previously studied marker genes [34]. A wide range of abnormal expression patterns were found in Decitabine-treated worms (Figure 9). This includes a reduced number of segmental stripes of *Pdum-engrailed* in worms with few morphologically-visible segments (Figure 9A-A’’), reduced expression of *Pdum-dlx,* which is normally expressed at the basis of anal cirri and in parapodia, on one side of the worm (Figure 9B-B’’), as well as ectopic expression of *Pdum-Hox3, Pdum-cdx* and *Pdum-piwiB* in developing segments (Figure 9C-E’’), which are consistent with persistent defect in growth zone functionality.

**Figure 9:**
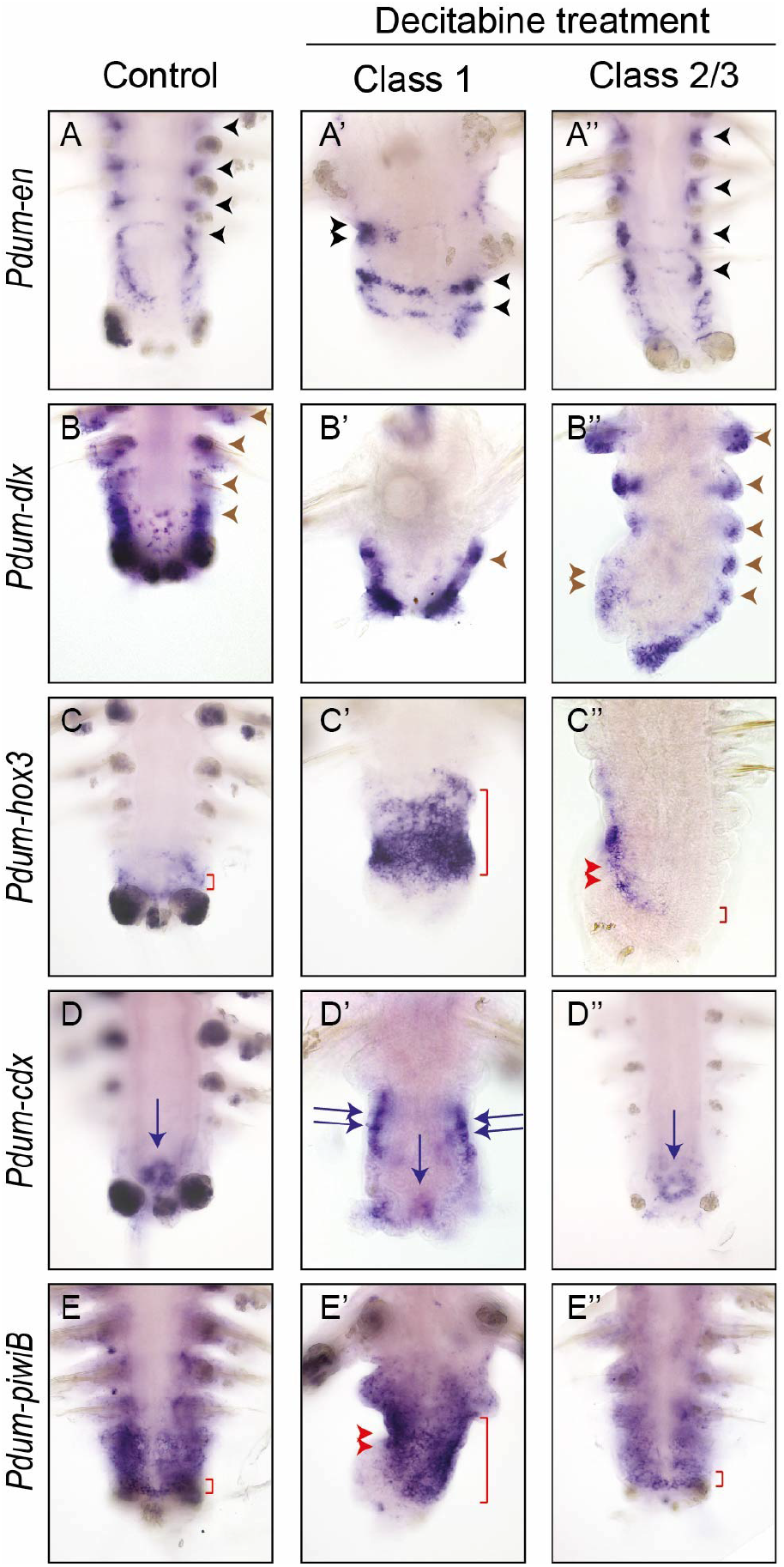
Gene expression at 25 days postamputation in worms that have been treated with Decitabine during regeneration. WMISH at 25 dpa of posterior parts of control worms and worms treated with 50μM Decitabine from 0 to 5dpa for selected markers, *i.e., Pdum-engrailed (Pdum-en;* segmental stripes), *Pdum-dlx* (parapodia and anal cirri) *Pdum-hox3* (growth zone), *Pdum-caudal/cdx* (pygidium and growth zone), and *Pdum-piwiB* (growth zone and segmental progenitors). Two Decitabine-treated worms with more or less altered morphologies (class 1 and class2/3) are shown for each gene. In A-A’’, black arrowheads point to segmental stripes of *Pdum-en* expression. In A’, double black arrowhead points to an incomplete expression stripe. In B-B’’, brown arrowheads point to expression of *Pdum-dlx* in developing parapodia. In B’’, reduced *Pdum-dlx* expression is observed on one side of the worm (double brown arrowhead). In C, red bracket delineates expression of *Pdum-hox3* in ectodermal growth zone. A very large domain of *Pdum-hox3* is found in class 1 worms (red bracket in C’). Abnormal expression pattern is also observed in class2/3 worms (red bracket and double red arrowhead in C’’). In D-D”’, blue arrows point to expression of *Pdum-cdx* in posterior gut region and double blue arrows to abnormal expression of the gene in developing segments (D’). In E-E’’’, red bracket delineates strong expression of *Pdum-piwiB* in mesodermal growth zone. The gene is also expressed in the developing mesoderm in a graded manner. In E’, strong and large expression is found throughout the regenerated region (red bracket and double red arrowhead).

Finally, we investigated whether Decitabine might have effects on a still longer time period. We treated worms with Decitabine from 0 to 5dpa, then put them in normal sea water until 25dpa, performed a second amputation one segment anterior the first amputation plan (meaning that we eliminated the regenerated region plus one segment) and scored these worms at several time points until 18 days post second amputation (18dpSa; Additional file 12: Figure S10A). Control and Decitabine-treated worms regenerated properly and similarly after this second amputation and were able to add new segments at a similar rate (Additional file 12: Figure S10B).

A slight but significant delay was however observed for worms treated with 10μM Decitabine at 18dpSa (Additional file 12: Figure S10B) and about 10-15% of Decitabine-treated worms showed minor defects in parapodia development (Additional file 12: Figure S10C). A same proportion of worms that were class 1 or 2 at 25dpa showed abnormalities after second amputation at 18dpSa (Additional file 12: Figure S10C). Multiple correlation analysis showed that, for both controls and Decitabine-treated worms, only positive correlations were found (Additional file 12: Figure S10D, E). Therefore, Decitabine treatment after a first amputation has only very minor effects on regeneration and post-regenerative posterior growth occurring after a second amputation.

Taken together, our data show that Decitabine decreases methylation level in *P. dumerilii* affecting larval development and regeneration. During regeneration it impairs post-regenerative posterior growth occurring in the absence of the drug, a long-term effect that could be due to defects in regeneration of the stem cell-containing growth zone.

## DISCUSSION

A wealth of studies, mainly conducted in mammals pointed out that cytosine DNA methylation modulate gene expression and is of primary importance for the regulation of embryonic development and stem cell properties [7]. Mechanisms and biological roles of this epigenetic modification in non-vertebrate species and in other processes such as regeneration, has however been much less studied. This is at least in part due to the fact that canonical non-vertebrate developmental models such as *Drosophila melanogaster* and *Caenorhabditis elegans* [60] and canonical regeneration models such as planarians [61], almost entirely lack 5mC and are therefore of no help for understanding its functions in nonvertebrates. In this article, we study cytosine DNA methylation and its roles during development, regeneration and post-regenerative growth in an emerging developmental and evolutionary biology model species, the marine annelid *Platynereis dumerilii.*

### P. dumerilii *genome displays a high and dynamic level of CpG methylation*

One of the main aims of our study was to define the extent and pattern of DNA methylation in the annelid *P. dumerilii.* Three main types of methylation patterns are classically described in animals: (i) global high-level methylation often presented as characteristic of vertebrates, in which a large majority of CpGs are methylated; (ii) mosaic low-/intermediate-level methylation in which only some genomic regions are methylated (interspaced with non-methylated ones) found in many diverse non-vertebrates; (iii) ultra-low/no methylation found in some species such as *D. melanogaster, C. elegans* and *Schmidtea mediterranea* [10,44,46,62]. Here, we combined *in silico* (computation of CpG o/e ratios [40,44]) (Figure 1A) and experimental (genomic DNA digestion with methylation-sensitive enzymes and LUminometric Methylation Assay (LUMA [49,50])) (Figure 1B, C) approaches to point out high levels of CpG methylation in *P. dumerilii* (up to more than 80% at some developmental stages) comparable to those of mammalian somatic cells [5,63]. While this high-level vertebrate-like methylation may seem surprising in a non-vertebrate species, it has also been demonstrated by whole-genome bisulfite sequencing in the sponge *Amphimedon queenslandica* (and is supported by the low CpG o/e ratios of this species) [64], showing that high-level methylation can also occur in non-vertebrate species. While our data indicate gene body methylation, as CpG o/e ratio calculation (Figure 1A) and bisulfite pyrosequencing (Figure 1D) were done on coding regions, further analyses, in particular whole-genome bisulfite sequencing, will be required to better characterize *P. dumerilii*genome methylation, for example to define whether there is 5mC depletion at gene promoters, like in *A. queenslandica* and vertebrates [64]. In addition, the question whether high-level genome methylation could be more widespread in animals than expected, deserves to be experimentally addressed, as patterns and extent of DNA methylation are known in rather few animal species so far and CpG o/e ratio calculations suggested that high-level methylation might exist in other non-vertebrate animals [44] (Figure 2).

An intriguing observation made about *P. dumerilii* DNA methylation is the sharp decrease of 5mC level, observed by LUMA, from more than 80% of methylated CCGG sites at embryonic/larval stage (12 to 72 hours post-fertilization, hpf) to about 60-65% at post-larval stages (5 and 15 days post-fertilization, dpf) (Figure 1C). It is important to note that we were unable for technical reasons, to obtain LUMA data from very early developmental stages, the earliest studied one being 12hpf which roughly corresponds to a late gastrula stage [32]. Later time points, 24, 48 and 72hpf, correspond to organogenesis stages during which many cells and tissues are differentiating. We cannot extrapolate methylation levels of earlier 0-12hpf stages, neither can we exclude that demethylation after fertilization may occur followed by remethylation during cleavage/early gastrulation stage, similarly to what has been observed in mammals [7]. In mammals, methylation level remains high in somatic cells from epiblast stage onwards and a transient decrease of this level only occurs in the germ cell lineage [7]. By contrast in *P. dumerilii,* a sharp decrease in DNA methylation level happens in the 72hpf to 5 and 15dpf time period, which corresponds to a major transition in the life cycle of the animal, the metamorphosis of the larva into a benthic feeding juvenile worm that adds new segments by posterior growth [32,33]. Such changes of DNA methylation levels at critical developmental transitions have also been reported in other non-vertebrates, for example at the time of metamorphosis in the oyster *Crassostrea gigas* [30] and in *Xenopus* [65], or during cast attribution in social hymenopterans *(e.g.,* [38,66]). Changes of DNA methylation level could therefore be important for key developmental transitions in distantly related animals, a tempting hypothesis that nevertheless has to be experimentally tested.

### P. dumerilii *possesses an ancestral-like repertoire of DNA methylation and NuRD toolkit genes that show dynamic expressions during development and regeneration*

We retrieved and analyzed a large dataset of genes encoding putative writers, modifiers, and readers of 5mC, as well as NuRD members and related proteins. This dataset corresponds to 17 gene families/subfamilies for which we reported the presence/absence and number of members in each of the 54 studied species (Figure 2). Using these data, we were able to infer parsimoniously the likely presence or absence of each gene family at key nodes of the metazoan tree (Figure 3). Our conclusions are in agreement with, and reinforce, those of previous studies made on a more limited set of gene families and/or studied species (*e.g.*, [22,26,29,64,67]). A complex DNA methylation and NuRD toolkit was present in the last common ancestor of all animals which owned at least one member of at least 16 of the 17 analyzed families/subfamilies, while the last common ancestor of eumetazoans and bilaterians possessed at least one member of all 17 families (Figure 3). This toolkit has been remarkably conserved during animal evolution as shown by the small number of missing orthologs in most species (Figure 2), a number likely to be overestimated as most species only benefit from draft genome assemblies. Exceptions are species in which gene losses occurred frequently such as nematodes, some insects and flatworms. In these species, very low/no genome methylation was reported and several genes of the methylation toolkit (mainly *Dnmt* and *Uhrf* genes) are absent. Similar reduced toolkits are found in the placozoan *Trichoplax adhaerens* and the tardigrade *Hypsibius dujardini,* suggesting that these two species may also have no or very reduced genome methylation (as also hinted by CpG o/e values). Besides these extreme cases, there is no clear connection between the number of genes encoding 5mC machinery proteins and the pattern of genome methylation, inferred from CpG o/e calculation (Figure 2) or experimental analysis [64].

We identified members of all 17 gene families/subfamilies in *P. dumerilii* with singlecopy members for 16 of them (Figure 2; the only exception is the CHD3/4/5 subfamily for which four members were found), consistent with the hypothesis that *P. dumerilii* belongs to a slow-evolving lineage in which few gene loss and duplication events occurred [68,69]. To have a first glimpse of possible function of these genes in *P. dumerilii,* we looked at their expression at the mRNA level, using transcriptomic data (for developmental and adult stages) and a combination of transcriptomic and whole-mount *in situ* hybridization data for regeneration. Most DNA methylation and NuRD genes are expressed at most or all stages of *P. dumerilii* life cycle and during regeneration (Figures 4 and 5), which is expected for genes encoding epigenetic regulators likely involved in multiple steps of the life cycle of the animal. Many genes however show dynamic expression during embryonic/larval development, at juvenile and adult stages, as well as during regeneration. Interestingly, at the transition between larval/post-larval stages, expression of *Pdum-dnmt1* and *Pdum-uhrf,* which could be involved in DNA methylation maintenance like their vertebrate orthologs [16], decreases, while expression of *Pdum-tet* [18,19], likely involved in demethylation, increases (Figure 4), suggesting that the marked decrease in DNA methylation level observed at this transition (see above), could be due to either passive (Dnmt1-mediated) or active (Tet-mediated) demethylation, or to both mechanisms.

We also characterized expression of several DNA methylation and NuRD genes during regeneration and found these genes to be expressed at all stages of the process (Figures 4 and 5). Strikingly all the studied genes are expressed at stage 1 in cells of the wound epithelium and/or cells of the immediately adjacent segment, which would fit with an early role of these genes during regeneration. From stage 2 onwards, all genes are expressed in the regeneration blastema and subsequently in the regenerating segments, in patterns that are reminiscent of those previously reported for stem cell and proliferation genes [34], suggesting an expression in proliferating cells and therefore that DNA methylation could be important for the formation of regenerated structures from blastemal cells. These expression data are also consistent with the hypothesis that DNA methylation might be involved during multiple steps of regeneration in *P. dumerilii*.

### *Decitabine treatments suggest that DNA methylation is involved in* P. dumerilii *development, regeneration and post-regenerative growth*

To study the putative functions of DNA 5mC methylation during *P. dumerilii* development and regeneration, we treated larvae or regenerating worms with Decitabine (5-aza-2’-deoxycytidine), a hypomethylating agent which causes cell-division-dependent DNA demethylation by blocking Dnmt1 activity [56–58]. While widely used in vertebrates, especially humans, this chemical has only been sparsely used in non-vertebrates [70–73] and, to our knowledge, never used during regeneration or in annelids. We therefore obtained evidence that Decitabine does indeed lead to DNA demethylation in *P. dumerilii* by showing a 2.5-fold decrease of CCGG methylation in 72hpf larvae treated with Decitabine for two days, as compared to control animals (Figure 6B). Decitabine is thus an appropriate tool to study possible roles of DNA methylation in *P. dumerilii.*

We observed three main effects for Decitabine. First, when applied on larvae from 1dpf to 3dpf, Decitabine produced important morphological defects but did not block development or led to larval death (Figure 6C). Some developmental processes, such as appendage (parapodia) and pygidium formation, are strongly affected (as shown by the extreme reduction of these structures in Decitabine-treated larvae), while others, such as segmentation, are less or not affected. This could point out differential requirements of DNA methylation for specific developmental process. Alternatively, it could be due to the mode of action of Decitabine which induces a progressive loss of 5mC through cell divisions, meaning that a significant decrease in 5mC level is probably achieved only several hours after the start of Decitabine treatment. In this view, Decitabine treatment may not affect developmental processes that happens before or soon after 24hpf, for example segmentation (segmental stripes of *Pdum-engrailed* expression can already be seen at 18hpf [74]), while strongly affecting processes starting later in development, for example pygidium formation which is still ongoing at 3dpf [75]. In the future, it would be interesting to further assess the effects of Decitabine on *P. dumerilii* development, in particular through the use of other temporal windows of treatment, including windows spanning early development (0-24hpf) which could produce much more severe defects, as observed in the oyster *C. gigas* in which gastrulation was severely impacted when Decitabine was applied on early development stages [71]. Effects of Decitabine on germ cell specification and development, which are well characterized in *P. dumerilii* [76], would also be an interesting topic for future examination, given the importance of DNA (de)methylation in the germ cell lineage [7].

A second clear effect of Decitabine is that it strongly delays regeneration when applied on worms for five days, from 0 to 5dpa (Figure 7A, B). Indeed, at day five after amputation, Decitabine-treated worms mostly reached stage 2 or 3, while controls worms reached stage 4 or 5. This is similar to what has been observed when applying, the proliferation inhibitor hydroxyurea on regenerating worms [34]. On one hand, this is consistent with the antiproliferative effect of Decitabine observed in humans in which this drug has been used for a long time, at high concentration, as an anticancer cytotoxic drug preventing DNA replication, thereby blocking cell proliferation and leading to cell death [77,78]. On the other hand, several elements suggest that the effect of Decitabine on regeneration in *P. dumerilii* could be due to demethylation and not to an impairment of DNA replication. The first and most compelling one is that we used low Decitabine concentrations that effectively led to significant demethylation (Figure 6B). As this Decitabine-induced demethylation can only occur after several cell divisions, it could not be obtained if Decitabine was used at concentrations that block cell divisions and has cytotoxic effect [77,78]. In humans, Decitabine is now used at low dosage to maximize its hypomethylating action, which elicits better anticancer responses than when used at higher concentrations [78]. Second, while there are many cell divisions which happens during larval development (1 to 3dpf time period; *e.g.,* [79]), Decitabine treatment leads to 3dpf larvae with a size and a body shape similar to control ones strongly arguing against an antiproliferative effect of Decitabine at the used concentrations. Third, after washing out the drug, Decitabine-treated worms were able to rapidly resume regeneration in a normal manner, which does not support significant cytotoxic effects of Decitabine. Our current hypothesis is therefore that Decitabine delays regeneration through its hypomethylating effect and therefore that DNA methylation may be required for proper regeneration in *P. dumerilii*. This hypothesis is at odds with what has been described in vertebrates in which, in contrast, demethylation has been suggested to be a driver of regeneration, in the axolotl limb [37], zebrafish fin [36] and the chick retina [80]. Additional experiments will be required to further test the requirement of DNA methylation in *P. dumerilii* regeneration and identify the involved mechanisms. An obvious possibility would be that massive demethylation induced by Decitabine may interfere with the expression of genes involved in regeneration, as suggested by reduced expression of *Pdum-Hox3, Pdum-piwiB* and *Pdum-engrailed,* in Decitabine-treated worms (Figure 7C), thereby hampering successful regeneration.

A third compelling effect of Decitabine is that, when applied during five days after amputation, it interfered with post-regenerative posterior growth that occurs once regeneration has resumed and been completed in the absence of the drug (Figure 8A, B). These worms showed a reduced rate of segment addition compared to control animals and displayed segments with various abnormalities and severely affected gene expression patterns at 25dpa, *i.e.,* 20 days after drug removal (Figures 8 and 9). Based on multiple correlation analysis (Additional file 11: Figure S9) and altered gene expressions of stem cell, growth zone and segmental markers in Decitabine-treated worms at 5dpa (Figure 7C), we suggest that drug-mediated DNA hypomethylation affects gene expression in stem cells of the growth zone, thereby impairing its functionality and subsequent posterior growth. Abnormal expressions of growth zone markers are still observed in some worms at 25dpa (Figure 9), consistent with the hypothesis of an epigenetic modification (hypomethylation) that has been transmitted during the many cell divisions that occur during the 5dpa to 25dpa time period. Along the same line, parapodia formation, which starts after the drug has been removed, is also strongly affected in many Decitabine-treated worms, suggesting that DNA hypomethylation might have been transmitted from growth zone stem cells to parapodial progenitors, leading to altered gene expression during parapodia formation, as shown for *Pdum-dlx* (Figure 9), and defects in this process.

Our data therefore suggest that 5mC DNA methylation is important for the function of stem cells of the growth zone in *P. dumerilii, i.e.,* somatic adult stem cells, and their capability to produce differentiated structures such as segments and appendages. Growth zone stem cells express a set of genes, such as *piwi, vasa* and *nanos,* which constitutes the so-called Germline Multipotency Program (GMP) shared with primordial germ cells and pluripotent/multipotent somatic stem cells in other animals [33,81]. In mammals, in the absence of DNA methylation, ES cells keep their stem cell identity and their self-renewal ability, but their differentiation is almost completely abolished, due to failure in upregulating germ-layer-specific genes and silencing pluripotency genes (*e.g.*, [82]). Roles of DNA methylation have also been described in mammalian adult stem cells, for both stem cell selfrenewal and differentiation capabilities *(e.g.,* [83–85]). A tempting hypothesis is therefore that Decitabine-induced hypomethylation in *P. dumerilii* growth zone stems cells may alter their ability to produce differentiated cell lineages required for segment and appendage formation. In agreement, ectopic expressions of growth zone markers, *Pdum-Hox3, Pdum-cdx* and *Pdum-piwiB* were observed in developing segments of 25dpf worms that have been treated from 0 to 5dpf with Decitabine (Figure 9), suggesting that silencing of stem cell markers in segmental progenitors/differentiated cells might not be properly done in a hypomethylated context. Further test of this hypothesis would require to specifically assess methylation levels in growth zone stem cells and their progeny, which is currently not possible due to the lack of tools to isolate and culture these cells.

### Conclusions

We provide data that strongly suggest that the genome of *P. dumerilii* is highly methylated and that methylation level changes during development. We show that *P. dumerilii* harbors a mostly single-copy repertoire of DNA methylation and NuRD genes that show dynamic expressions during development and regeneration. Using the hypomethylating drug Decitabine, we obtained functional data in favor of an involvement of 5mC DNA methylation during development, regeneration and post-regenerative growth. These treatments strongly suggest in particular that Decitabine-induced hypomethylation of stem cells of the growth zone alters their capability to produce differentiated segments. Taken together our data provide the first evidence, outside vertebrates, of roles of 5mC epigenetic mark during regeneration and in stem cells involved in growth. Our study also lays the groundwork for using *P. dumerilii* as a new non-vertebrate model to study 5mC DNA methylation, and other epigenetic regulations, in particular those involving histone modifications, during development, regeneration and stem cell-based growth.

## MATERIALS & METHODS

### P. dumerilii *gene identification and cloning*

Putative *P. dumerilii* orthologs of genes known to encode protein involved in 5mC methylation or to belong to NuRD complex (and related proteins) were identified by BLAST searches [86] on available transcriptomic and genomic data, using *H. sapiens* and *M. musculus* protein sequences as queries. Sequences of all identified genes can be found in Additional file 4. Conserved domains in *P. dumerilii* and *H. sapiens* proteins (Additional file 3: Figure S2) have been identified using InterPro 80.0 [87] and the NCBI online software CD-Search Tools [88]. High-fidelity PCR with gene-specific primers (listed in Additional file 1: Table S6) was used to amplify gene fragments using as template cDNAs from mixed larval and regenerating stages with, for some genes, a touchdown approach. PCR products were purified (740609, Macherey Nagel), and TA cloned in the pCR2.1 vector (K450001, ThermoFisher), following manufacturers’ instructions. Sequences of cloned genes were verified by Sanger sequencing (Eurofins Genomics). GenBank accession numbers for *P. dumerilli* genes: MW250929 to MW250948.

### Phylogenetic analyses and establishment of orthology relationships

Putative members of all studied 5mC machinery gene families were identified by sequence similarity searches (reciprocal best BLAST hit approach) on publicly available genome sequences of 51 different species belonging to all major animal clades (Figure 2) using *H. sapiens* and *M. musculus* sequences as queries. Redundant sequences were manually discarded and, in some cases, incomplete short sequences were concatenated. All identified sequences can be found in Additional file 4. Databases used to retrieve these sequences are listed in Additional file 1: Table S7.

Multiple alignments were obtained using MUSCLE 3.8.31 [89] (available on the MPI Bioinformatics Toolkit platform [90]) and subsequently manually improved using SEAVIEW [91]. Maximum likelihood (ML) analyses were performed using the online available PHYML software [92,93]. Amino acid substitution model was defined by the software using SMS with the Akaike Information Criterion [94]. Other parameters (such as proportion of invariable sites and Gamma shape parameter) were estimated from datasets by the software. Tree improvement method was NNI and statistical supports for internal branches of trees was assessed by approximate Likelihood-Ratio Test (aLRT) [95]. Phylogenetic trees were handled using FigTree 1.4 (http://tree.bio.ed.ac.uk/software/figtree/).

### CpG o/e ratio calculations

Notos [40] was used to calculate and model CpG o/e (observed/expected) as described in Aliaga et al. [44] for *P. dumerilii* reference transcriptome and transcriptomes of some of the 54 studied species for which CpG o/e ratios were not previously determined [44]. The formula (CpG / (C * G)) * (L^2 / L-1) and a minimum length of 200 bp were used.

### P. dumerilii *breeding culture and posterior amputation procedure*

*P. dumerilii* embryos, larvae, and juvenile worms were obtained from a breeding culture established at the Institut Jacques Monod (Paris, France). For regeneration experiments, amputations of posterior parts were performed on juvenile worms of 3-4 months and 30 to 40 segments as previously described [34].

### *Study of DNA methylation levels in* P. dumerilii

Methylation of genomic DNA (gDNA) of *P. dumerilii, N. vectensis* (adult), *D. melanogaster* (adult), *B. floridae* (adult), *H. sapiens* (HCT116 cells), and *M. musculus* (embryonic stem cells, mESC) was assessed by digesting 500ng to 800ng of gDNA with 1μL of restriction enzymes HpaII or MspI in 50μL containing 1X FastDigest green buffer (FD0514 and FD0544, Thermo Scientific) for 15min at 37°C. 300ng of undigested gDNA and from each restriction reaction were loaded in a 1% agarose gel with molecular weight marker (N3232, NEB). 160V current was applied for 30min in a Midigel tank (370000, Apelex) and ethidium bromide was used with UV-transilluminator for revelation.

To evaluate global DNA methylation levels, we used LUminometric Methylation Assay (LUMA) based on enzymatic digestion and pyrosequencing [49]. Two or three fertilizations were used for each analyzed larval/post-larval stages, per replicate. Samples were washed twice with 200mM PBS 0.1% Tween on ice before freezing at −80°C. DNA was extracted using the DNA/RNA All prep kit (80204, Qiagen) following manufacturer instructions. DNA was stored at −20°C prior to LUMA analysis. Prior to LUMA analysis, DNA integrity and sample purity were assessed using Tapestation (Agilent). Only samples with a DNA Integrity Number superior to 8 were kept for subsequence analysis. LUMA experiments were performed as in Karimi et al. [49] with internal controls (HCT116 WT and DKO (Dnmt1-/-; Dnmt3b-/-)) and technical duplicates. 250ng of gDNA were digested with HpaII+EcoRI or MspI+EcoRI for 4 hours at 37°C. Then, samples were analyzed in a PyroMark Q24 (Qiagen). The instrument was programmed to add dNTPs as following: dATP; dGTP + dCTP; dTTP; H2O as control; dGTP + dCTP; dATP; dTTP. Peak heights were calculated using the PyroMark Q24 software (Qiagen). The HpaII/EcoRI and MspI/EcoRI ratios were calculated as (dGTP + dCTP)/mean (dATP; dTTP) for the respective reactions. The percentage of methylated CCGG sites was defined as: 100 x [1-(HpaII/EcoRI)/(MspI/EcoRI)].

To measure CpG methylation levels in the *14-3-3-like* and *Histone H4* genes, we used bisulfite pyrosequencing. Pyrosequencing primers were designed using the PyroMark Assay Design Software 2.0 (Qiagen). 500ng of genomic DNA was subjected to bisulfite conversion using the EpiTect Bisulfite Kit (Qiagen, Catalog No. 59124). PCR reactions were performed in a final volume of 25 μL, using the Pyromark PCR kit (Qiagen, Catalog No. 978703), with one of the primers biotinylated and containing 12.5ng of bisulfite-treated DNA. The initial denaturation/activation step was performed at 95°C, 15 min, followed by 50 cycles of 30 sec at 94°C, 30 sec at 54°C, 45 sec at 72°C and a final extension step at 72°C for 10 min. The quality and the size of the PCR products were evaluated by running 5μL of each PCR product on 1.5% (w/v) agarose gel in a 0.5X TBE buffer. Biotinylated PCR products (20μL) were immobilized on Streptavidin-coated Sepharose beads (GE Healthcare, 17-5113-01). DNA strands were separated using the PyroMark Q24 Vacuum Workstation, biotinylated single strands were annealed with 0.375μM sequencing primer and used as a template for pyrosequencing. Pyrosequencing was performed using PyroMark Q24 Advanced (Qiagen, Catalog No. 9002270) according to the manufacturer’s instructions, and data about methylation at each CpG was extracted and analyzed using the PyroMark Q24 Advanced 3.0.0 software (Qiagen).

### *Whole-mount* in situ *hybridizations (WMISH) and imaging*

For probe synthesis, plasmid containing appropriate cDNA were purified (740588 and 740412, Macherey Nagel), digested (NEB enzyme) and used to produce digoxygenin-labeled RNA antisense probes (Synthesis with Roche reagents: 11093274910 and 10881767001 or 10810274001; Purification with 740955, Macherey Nagel) as previously described [33]. WMISH were done as previously described [34,96]. Bright field images were taken on a Leica microscope. Adjustment of brightness and contrast were performed using Photoshop software.

### Decitabine and RG108 treatments

Stock solution of 200mM Decitabine (5’-Aza-2’-deoxycytidine) and 10mM RG108 (N-Phthalyl-L-Tryptophan) in DMSO were diluted in sea water to obtain different concentrations as described in the Results section. DMSO controls correspond to a concentration of DMSO in sea water corresponding to that of 100μM Decitabine condition. In order to assess methylation levels with LUMA after Decitabine or RG108 treatment, larvae were kept 2 days (1 to 3dpf) in 30ml of Decitabine- or RG108-containing sea water before washing and freezing. For morphological studies, larvae were washed out, kept in normal sea water and fed from 5dpf onwards. Larvae were anesthetized with 7.5 % MgCl2 before observations.

Amputated worms were placed individually in 12 wells plate in 2ml of Decitabine solution or control solution that was changed every day for five days. For posterior growth analyses, worms were subsequently placed in normal sea water until 25dpa. For some experiments, at 25dpa, worms were amputated a second time one segment anterior to the first amputation plane, and posterior parts were fixed for WMISH.

### Scoring and statistical analysis

Scoring system established in Planques et al. [34] was used to score worms during posterior regeneration and post-regenerative posterior growth. Graphic representation of transcriptomic, LUMA and morphological experiments with corresponding statistical analyses were performed using Prism 7 software (GraphPad). Statistical test that have been used are indicated in legends of figures and supplementary figures. R was for used for multiple correlation computation and representation. Holm correction was applied for significance calculation [97].

## Supporting information

All supplementary figures

All supplementary tables

## ACKNOWLEDGEMENTS

We thank all Vervoort lab members for helpful discussions and feedback on the manuscript. We are grateful to Maxim Greenberg for helpful discussions and comments on the manuscript. We are grateful to the teams of Eric Rottinger, Sandra Duharcourt and Hector Escriva for providing us with genomic DNA from *N. vectensis, D. melanogaster* and *B. lanceolatum*, respectively. Animal facility members are thanked for their help with the worm culture. Tom Anerot-Rigo, Thibault Bidolet, Camille Kergavat, Maude Marchais, Erwan Martin, Edouard Riey and Anne Sauterau are thanked for their help with experiment set up. We acknowledge the Functional Epigenomics facility of the Epigenetics and Cell Fate unit (UMR7216) for its contribution to this work. We also thank Pierre-Antoine Defossez for helpful advices. This work was supported by funding from Labex ‘Who Am I?’ laboratory of excellence (No. ANR-11-LABX-0071) funded by the French Government through its ‘Investments for the Future’ program operated by the Agence Nationale de la Recherche under grant No. ANR-11-IDEX-0005-01, Centre National de la Recherche Scientifique, Université de Paris, Agence Nationale de la Recherche (grant TELOBLAST no. ANR-16-CE91-0007), the «Association pour la Recherche sur le Cancer » (grant PJA 20191209482), and the «Ligue Nationale Contre le Cancer» (grant RS20/75-20).

