## Supplementary material for "DNA methylation during development and regeneration of the annelid *Platynereis dumerilii*": All supplementary figures

### SUPPLEMENTARY MATERIALS

### SUPPLEMENTARY TABLES

**Additional file 1: Table S1: CpG o/e ratio obtained from Notos.**

**Additional file 1: Table S2: Sequences used for phylogenetic analyses.**

**Additional file 1: Table S3: Expression levels at different stages of regeneration based on RNA-seq data.**

**Additional file 1: Table S4: Lethality and autotomy induced by Decitabine treatment (short-term experiments).**

**Additional file 1: Table S5: Lethality and autotomy induced by Decitabine treatment (long-term experiments).**

**Additional file 1: Table S6: Primers used for *P. dumerilii* gene cloning and bisulfite pyrosequencing.**

**Additional file 1: Table S7: Databases used to retrieve sequences for phylogenetic analyses.**

### SUPPLEMENTARY DATA

**Additional file 4: List of all sequences used for phylogenetic analyses in fasta format.**

### SUPPLEMENTARY FIGURES

**Additional file 2: Figure S1: Additional CpG o/e ratio calculations.** Histograms of CpG o/e ratio for several species (whose name is indicated on top) for which this ratio has not been previously calculated. In each histogram, red line indicates the estimated density, vertical blue bar shows estimated mean value and shaded blue bar represents bootstrap confidence intervals of 95%. Clusters are those defined in Aliaga et al. [1]. Color code for metazoan groups is indicated and is as in Figure 2.

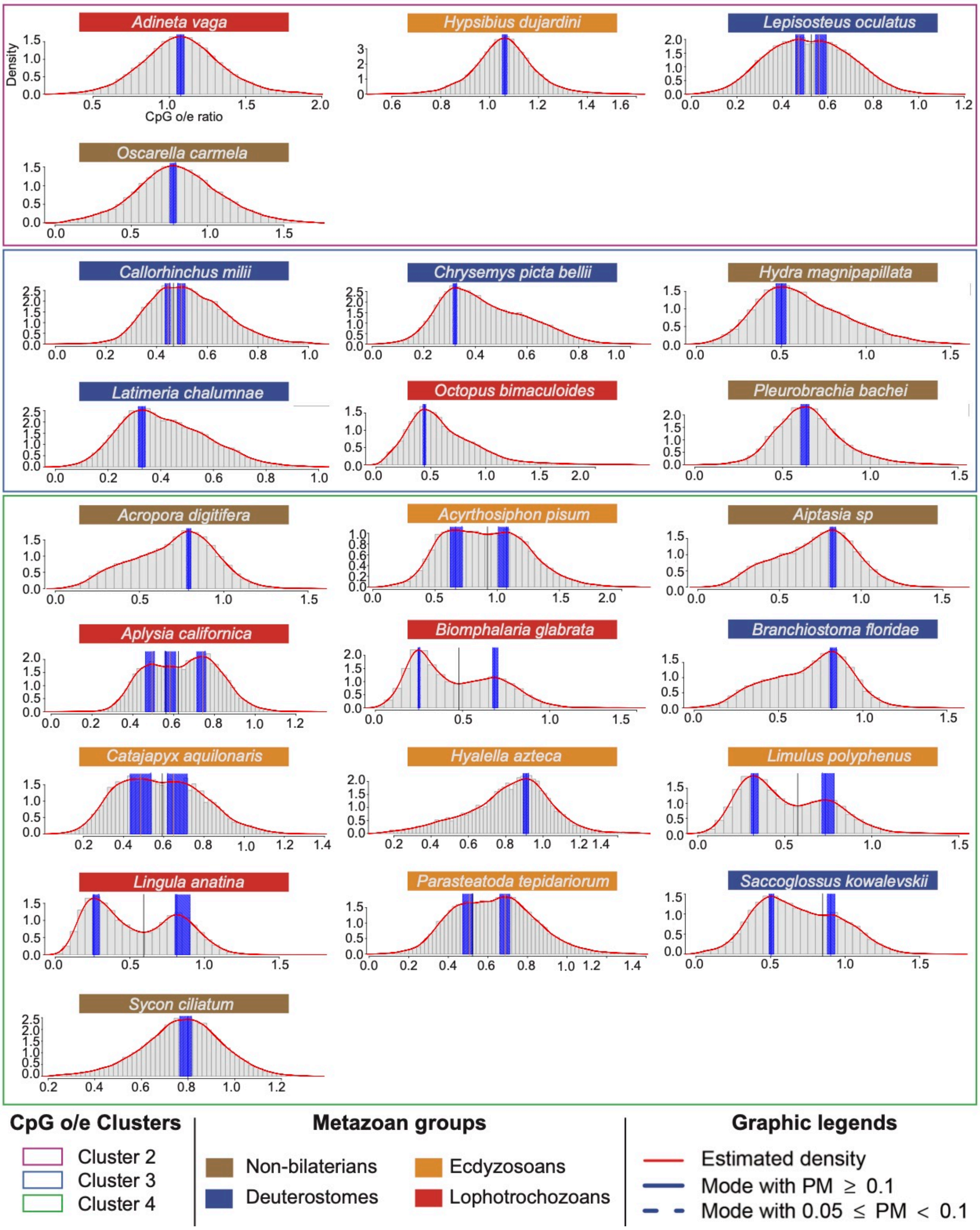

**Additional file 3: Figure S2: *P. dumerilii* 5mC and NuRD machinery genes and proteins.** All identified *P. dumerilii* genes are listed with the identification of the corresponding gene model in the Pdumbase reference transcriptome [2], excepted for *dnmt3*, identified in our unpublished regeneration transcriptome. Schematic representations of *P. dumerilii* (*Pdum*) and corresponding Human (*Hsap*) proteins are also shown, highlighting conserved domains found in these proteins and position of these domains.

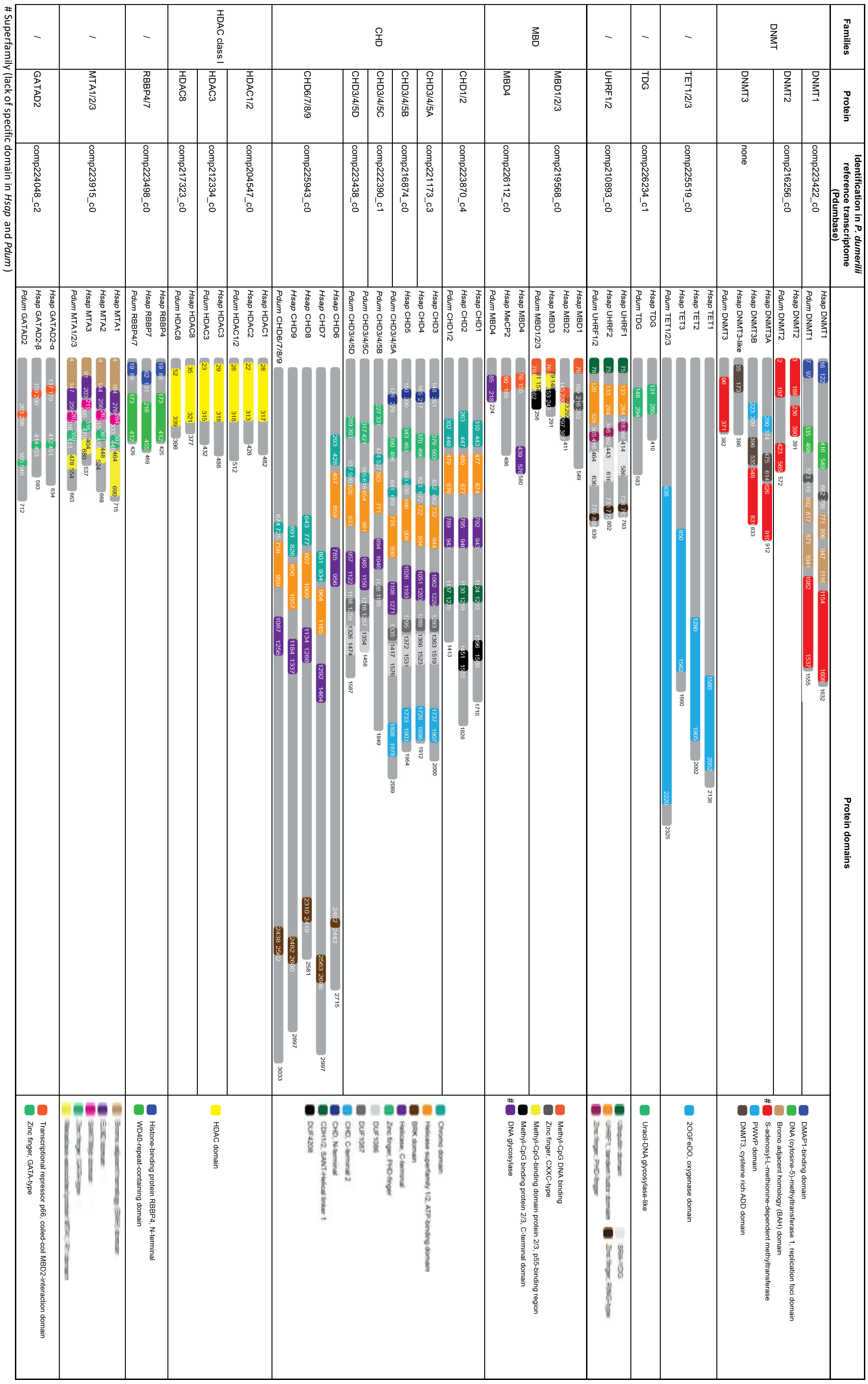

**Additional file 5: Figure S3: Phylogenetic trees of 5mC and NuRD machinery proteins.** Maximum-likelihood (ML) trees constructed with PhyML are shown. Statistical supports (aLRT values) for all nodes are indicated with a color code provided in the inset. Terminal branches are colored using the shown color code also used in Figure 2. *P. dumerilii* sequences are in bold and indicated by an arrow. (A) DNMT proteins. The three subfamilies DNMT1, 2 and 3 are indicated. In the DNMT3 subfamily, the vertebrate-specific groups DNMT3A, B and -like are also shown. (B) TET proteins. The three vertebrate-specific groups TET1, 2 and 3 are indicated. (C) TDG. (D) UHRF. The two vertebrate-specific groups UHRF1 and 2 are indicated. (E) MBD proteins. The two subfamilies MBD1/2/3 and MBD4/MeCP2 subfamilies are indicated. Vertebrate-specific groups are also shown. (F) CHD proteins. The three subfamilies CHD1/2, 3/4/5 and 6/7/8/9 are indicated. Vertebrate-specific groups are also shown. (G) Class I HDAC proteins. The three subfamilies HDAC1/2, 3 and 8 are indicated. Vertebrate-specific HDAC1 and 2 groups are also shown. (H) RBBP4/7 proteins. Vertebrate-specific RBBP4 and 7 groups are shown. (I) MTA1/2/3 proteins. Vertebrate-specific MTA1, 2 and 3 groups are shown. (J) GATAD2 proteins. Vertebrate-specific GATAD2-alpha and GATAD2-beta groups are shown.

### A. DNMT Proteins

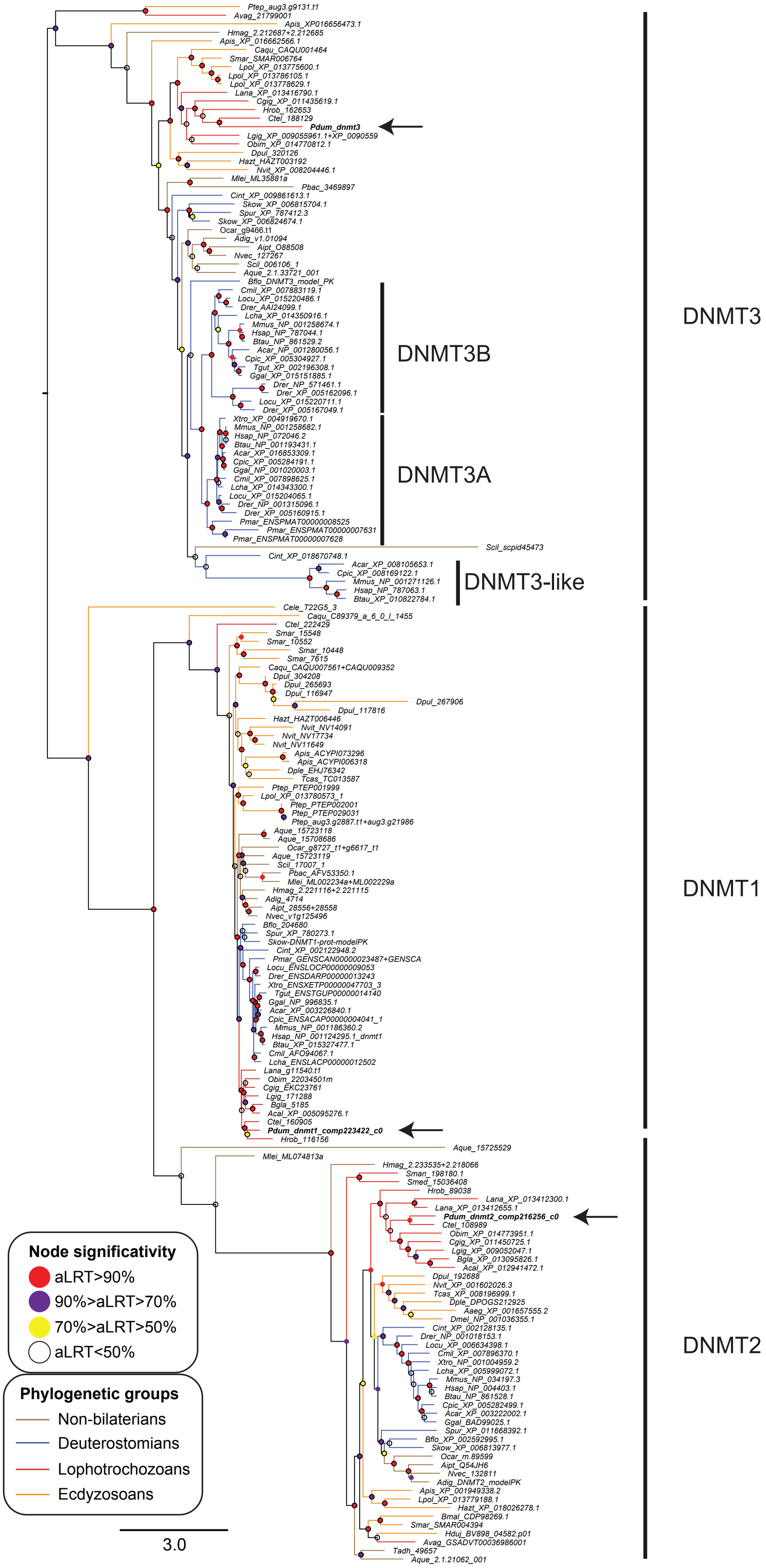

### B. TET Proteins

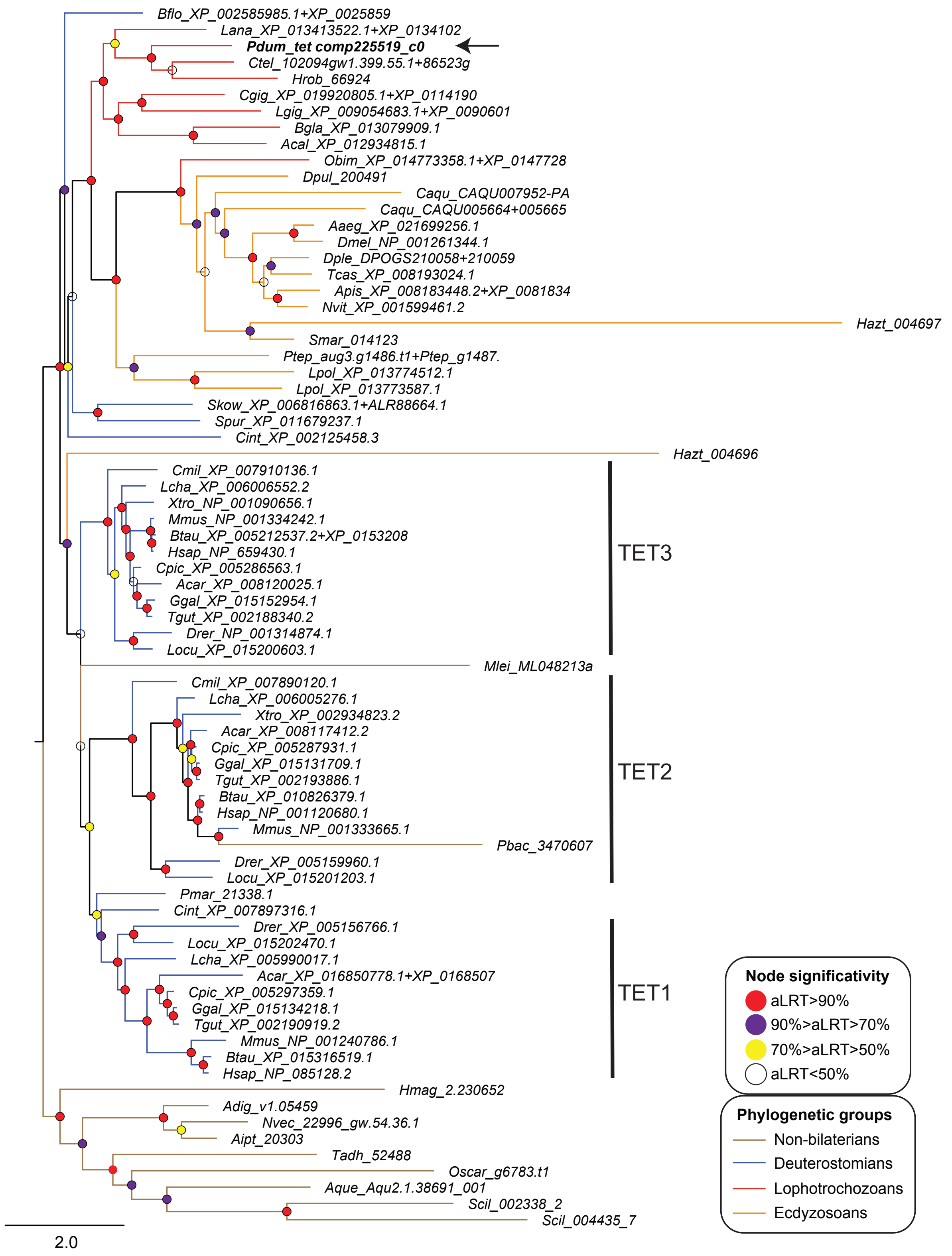

C. TDG Proteins

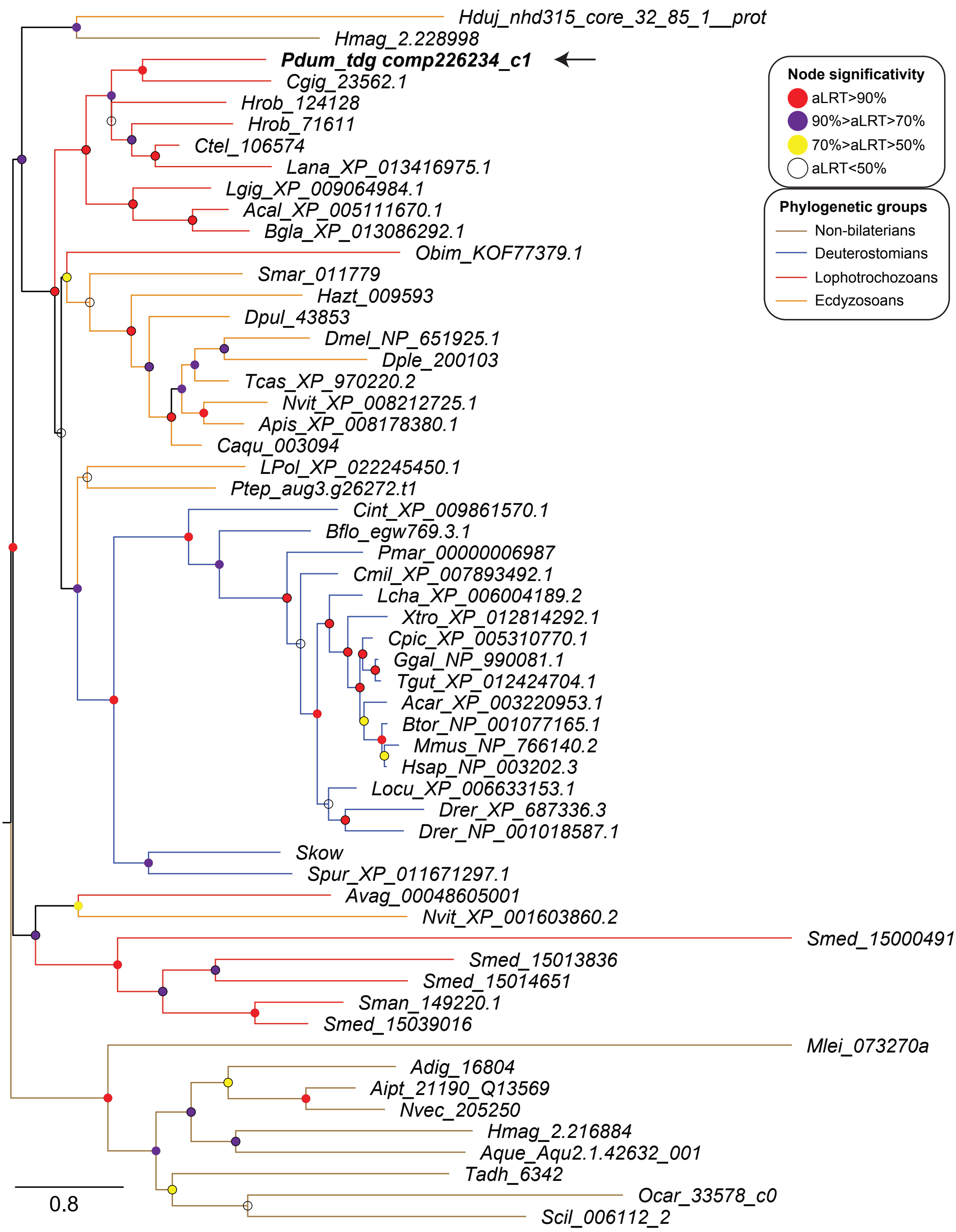

D. UHRF Proteins

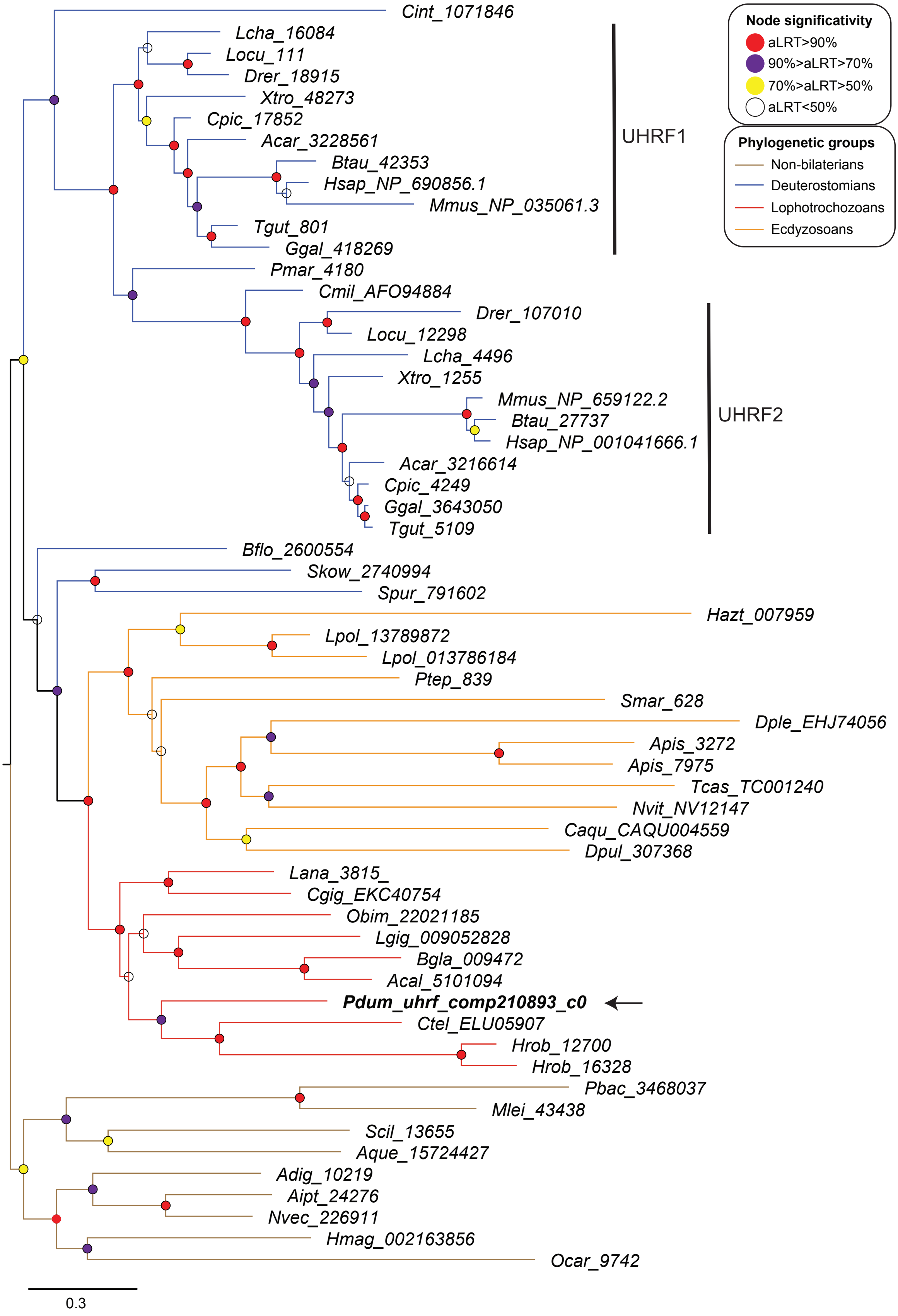

E. MBD Proteins

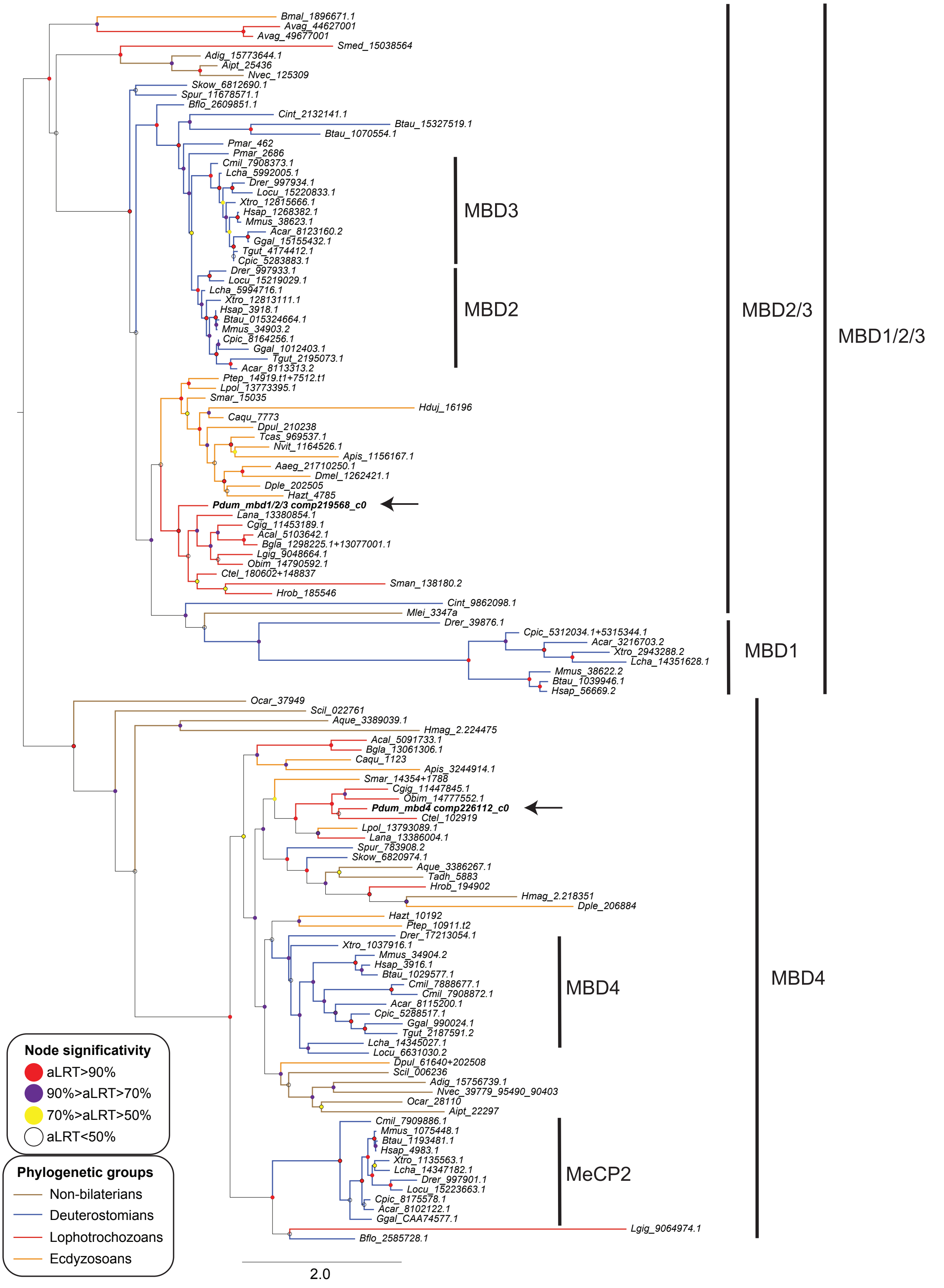

### F. CHD Proteins

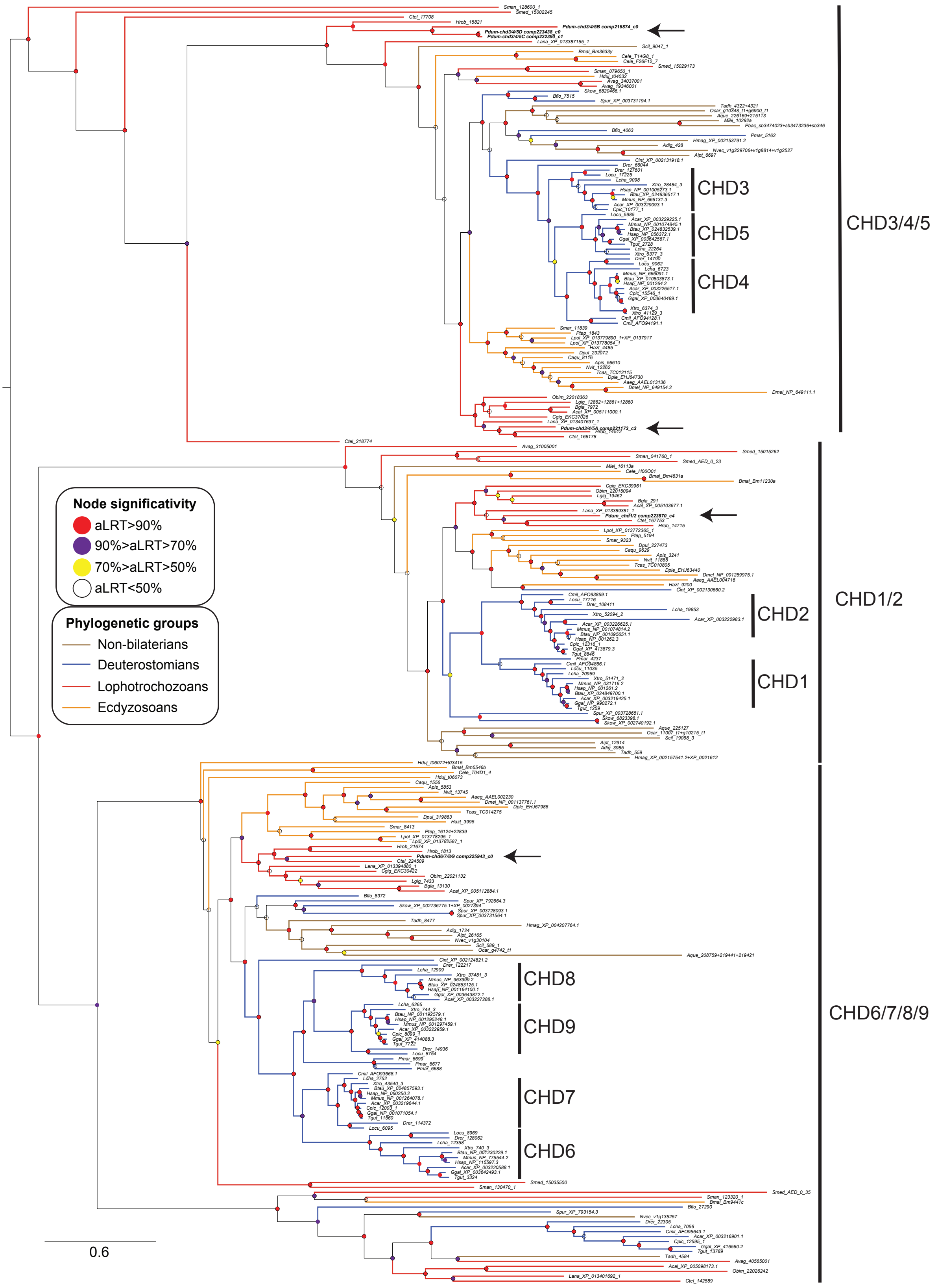

### G. HDAC Proteins

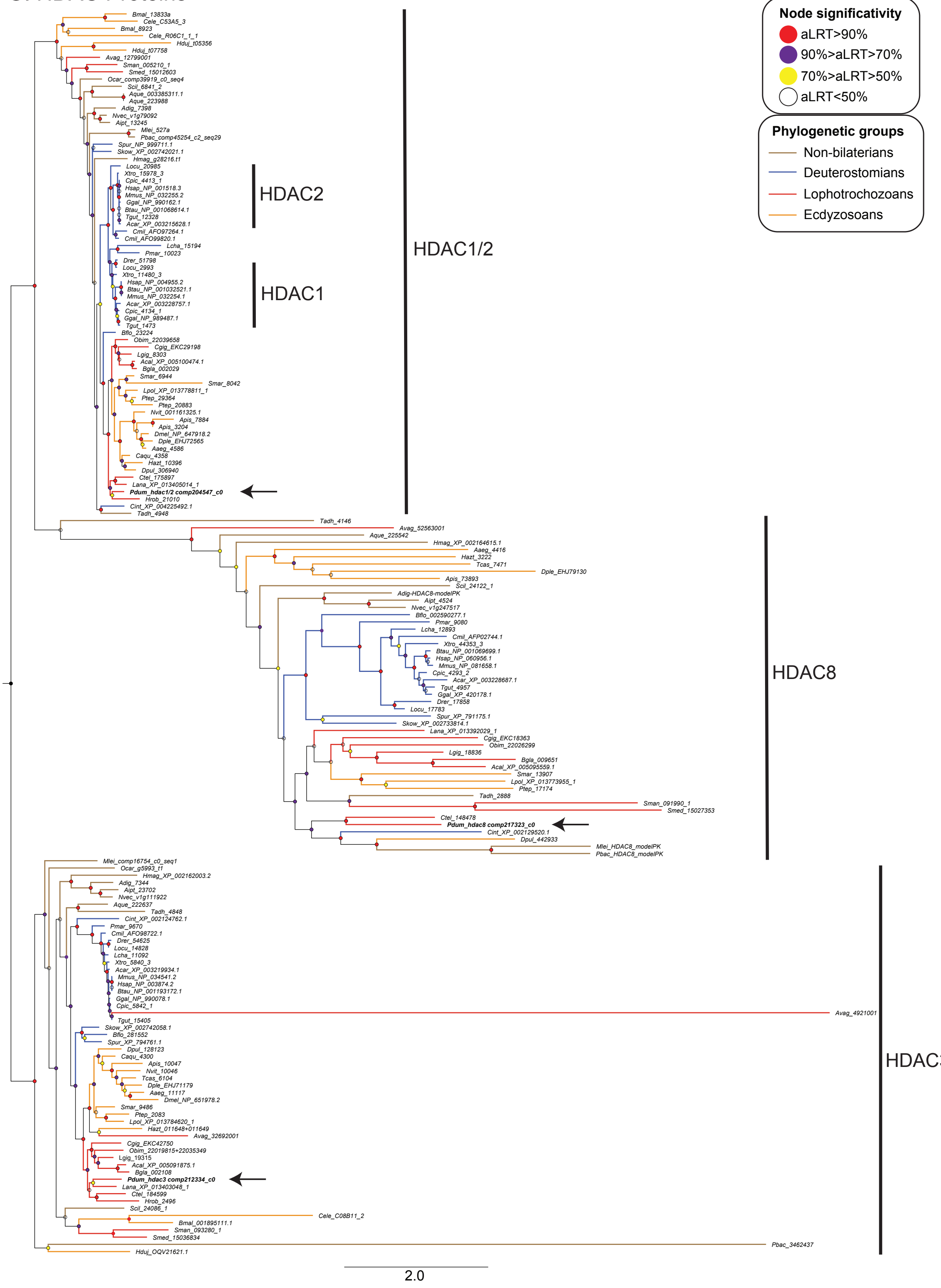

### H. RBBP4/7 Proteins

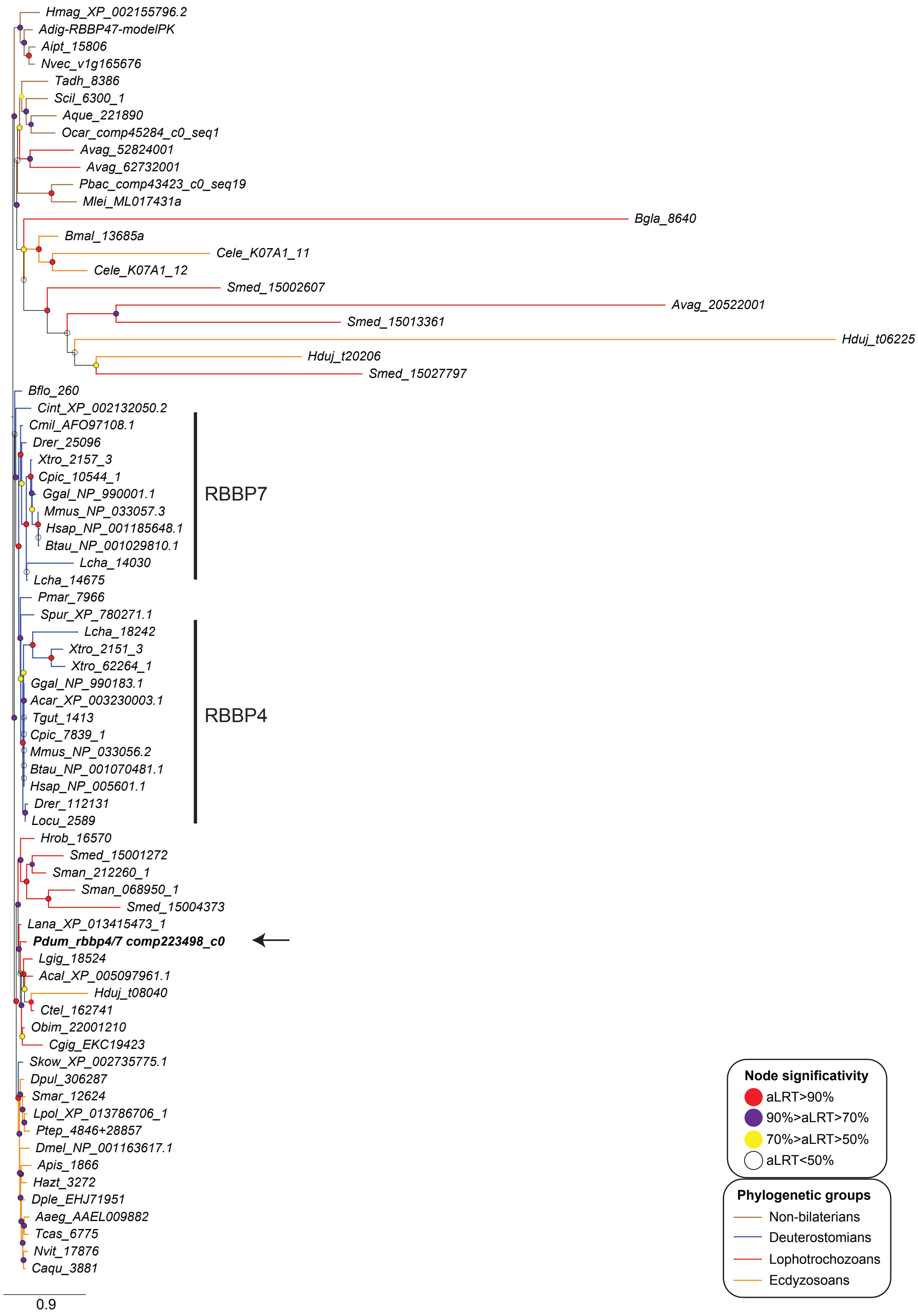

### I. MTA1/2/3 Proteins

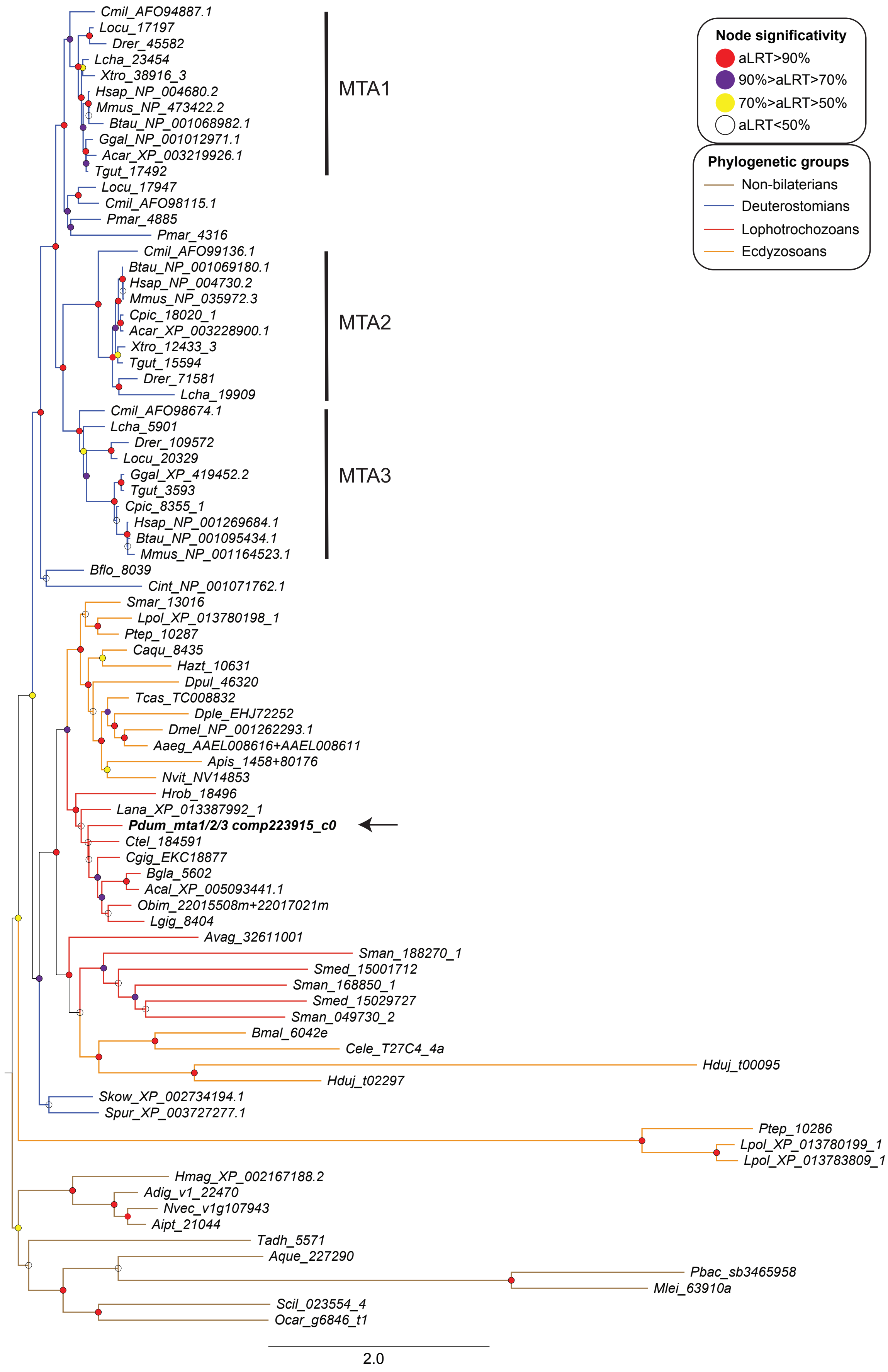

J. GATAD2 Proteins

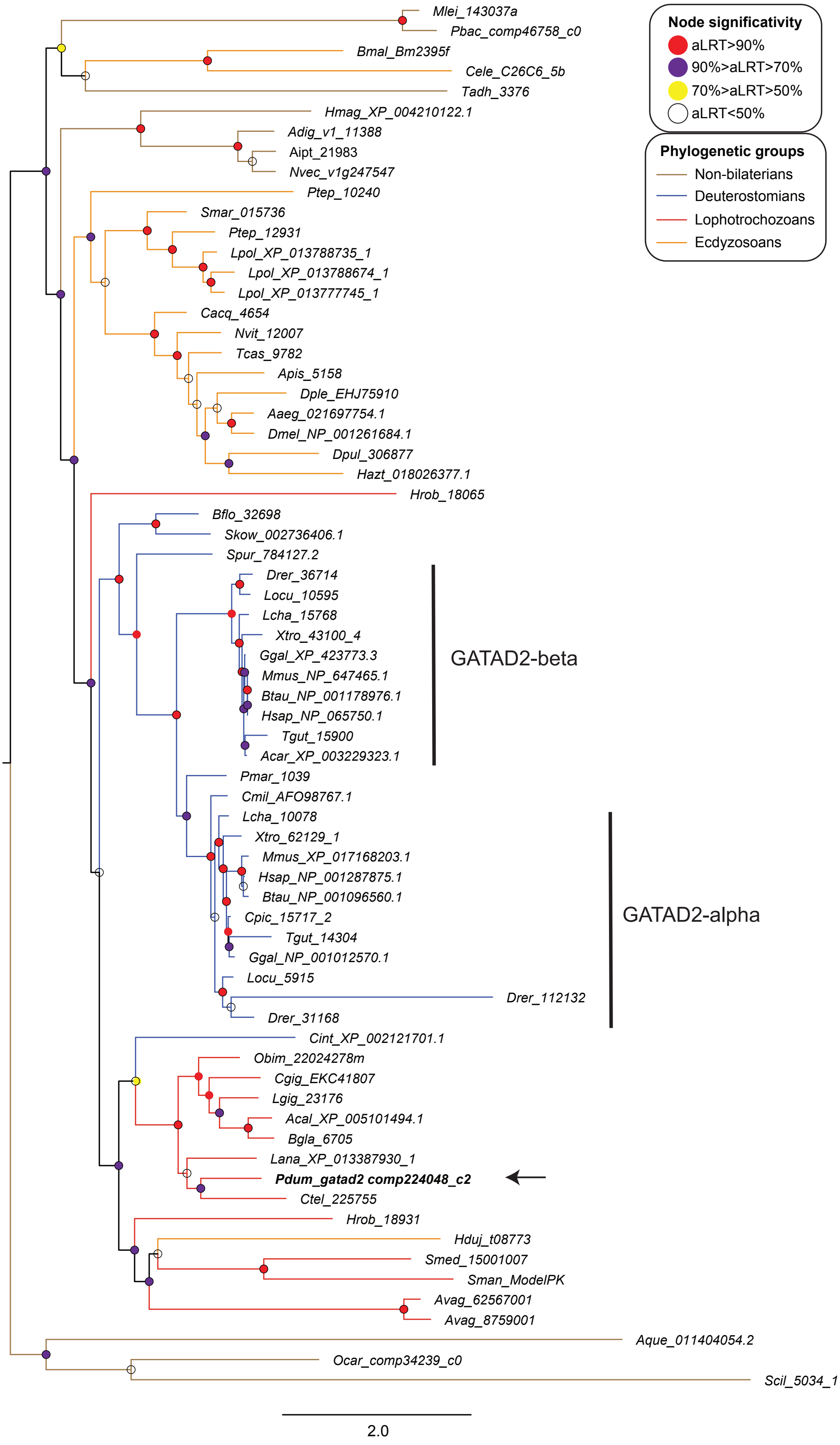

**Additional file 6: Figure S4: Expression level of 5mC and NuRD machinery genes during *P. dumerilii* development and along its life cycle.** Histogram reporting FPKM values at 19 stages (developmental and adult stages) indicated on the left for all indicated genes. Embryonic stages FPKM values and those for the other stages cannot be compared, as calculated from two independent RNA-seq studies. These two datasets are therefore separated by a black vertical dashed line. Black horizontal dotted line highlights a 5 FPKM threshold about which a gene can be considered as significantly expressed, as its expression can usually be detected by *in situ* hybridization [3]. Co-expression clusters are those defined by Chou et al. [3]. Hpf: hours post-fertilization; dpf: days post-fertilization; mpf: months post-fertilization.

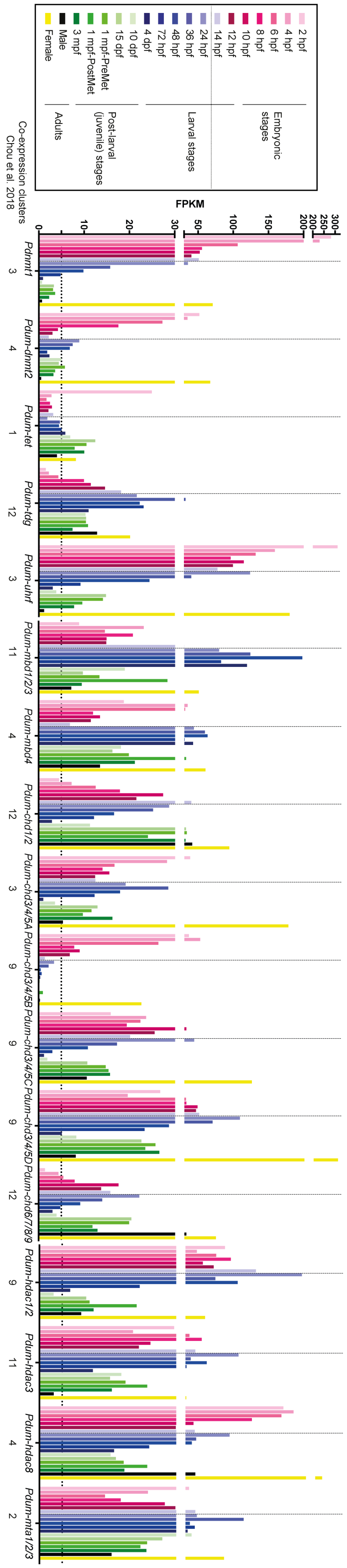
2 hpf
4 hpf
6 hpf
8 hpf
10 hpf
12 hpf
14 hpf
24 hpf
36 hpf
48 hpf
72 hpf
4 dpf
10 dpf
15 dpf
1 mpf-PreMet
1 mpf-PostMet
3 mpf
Male
Female
Embryonic stages
Larval stages
Post-larval (juvenile) stages
Adults
FPKM
Co-expression clusters Chou et al. 2018
Pdnmt1
Pdum-dnmt2
Pdum-tet
Pdum-tdg
Pdum-uhrf
Pdum-mbd1/2/3
Pdum-mbd4
Pdum-chd1/2
Pdum-chd3/4/5A
Pdum-chd3/4/5B
Pdum-chd3/4/5C
Pdum-chd3/4/5D
Pdum-chd6/7/8/9
Pdum-hdac1/2
Pdum-hdac3
Pdum-hdac8
Pdum-mta1/2/3

**Additional file 7: Figure S5: Additional expression data of 5mC and NuRD machinery genes during regeneration.** Whole-mount *in situ* hybridizations (WMISH) for the genes whose name is indicated are shown. In all panels, anterior is up and the regeneration stage for each picture is indicated. Dorsal (D) and posterior (P) views are shown. Light blue arrowheads = ectodermal growth zone, light blue arrows = ectoderm of developing segment, red arrows = groups of internal cells in the segment adjacent to the amputation plane.

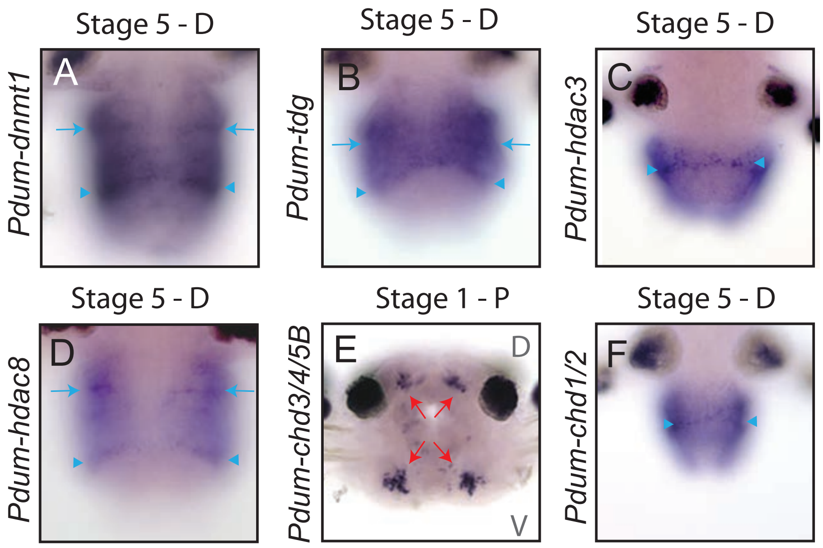

**Additional file 8: Figure S6: Schematic representation of expression patterns of 5mC and NuRD during regeneration.** In all panels, anterior is up and the regeneration stage for each schema is indicated. Ventral schematic representations are shown for all stages and, in addition, dorsal schematic representation is provided for stage 5. Color code for the different tissues is provided in the inset.

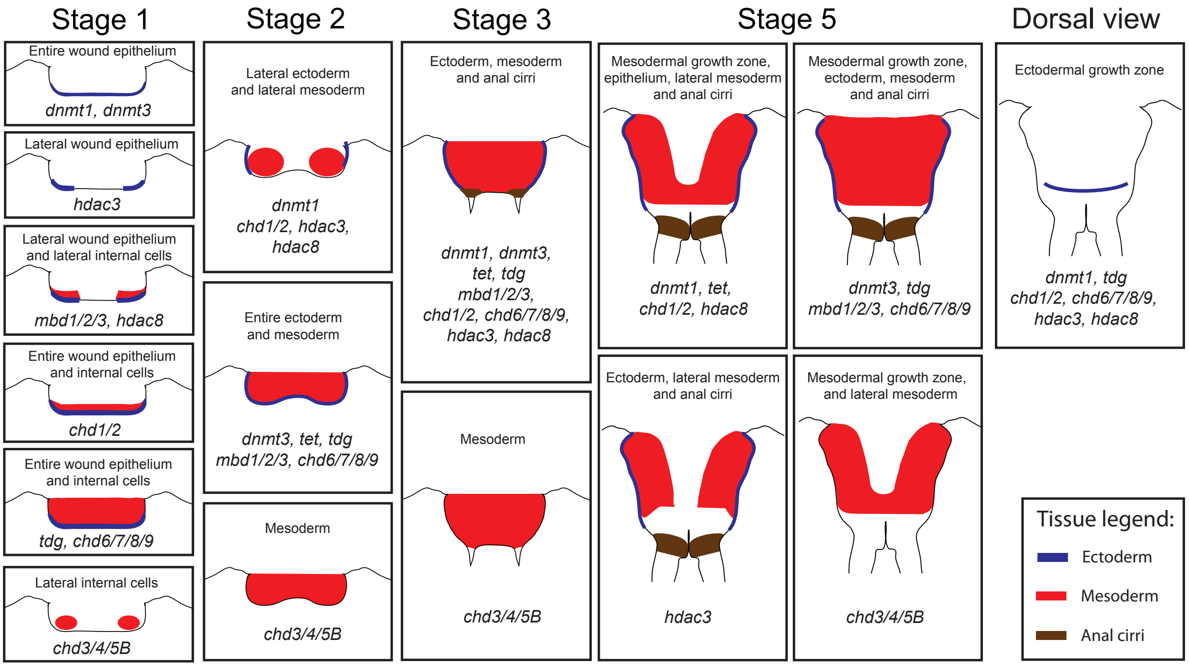

**Additional file 9: Figure S7: Rate of segment addition and individual scoring of Decitabine-treated worms.** (A) Graphic representation of rate of segment addition in controls (DMSO 0.01% and sea water) and Decitabine-treated worms (worms that showed autotomy are excluded). (B-F) Graphic representation of scoring of individual control (B and C) and Decitabine-treated (D-F) worms until 25dpa. Two experiments, mean ± SD. For rate of segment addition analysis, 1-way ANOVA was performed with Dunnett post hoc test (**: $p < 0.01$; ***: $p < 0.001$; ****: $p < 0.0001$). For conditions comparison, 2-way ANOVA was performed (Source of variation: Time $p < 0.0001$, Treatment $p < 0.0001$, Interaction $p = 0.0875$) with Dunnett post hoc test (*: $p < 0.05$; **: $p < 0.01$; ***: $p < 0.001$).

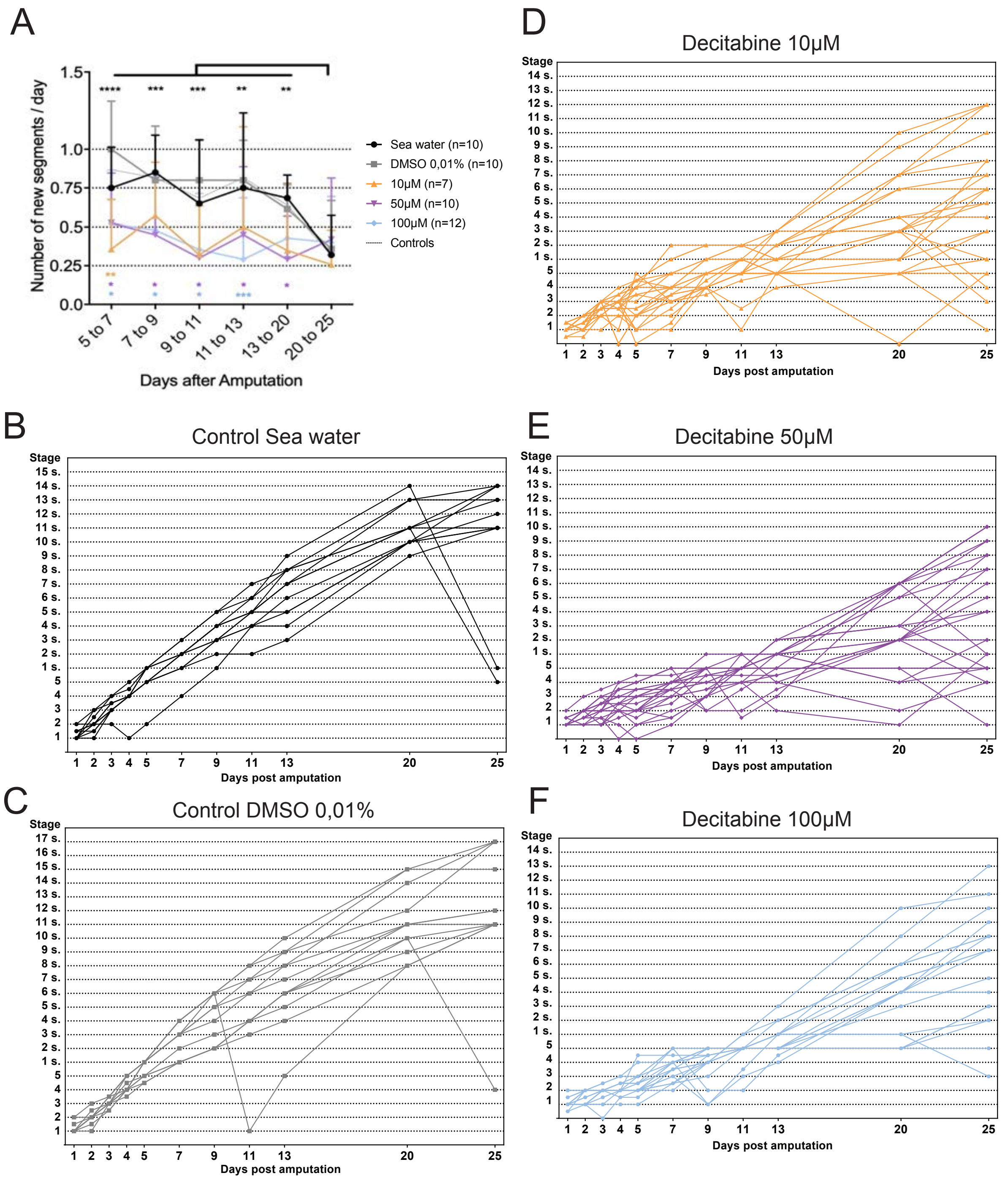

**Additional file 10: Figure S8: Classification of Decitabine-treated worms based on morphological defects.** Three classes of worms can be defined based on the reported morphological defects. For each class, representative worms at five different time points after amputation are shown. Green arrowheads = characteristic constriction between the non-regenerated and regenerated regions, pink arrows = very reduced or absent anal cirri, blue arrows = reduced/abnormal anal cirri, green arrows = narrowed regenerated region, black arrowheads = abnormal parapodia.

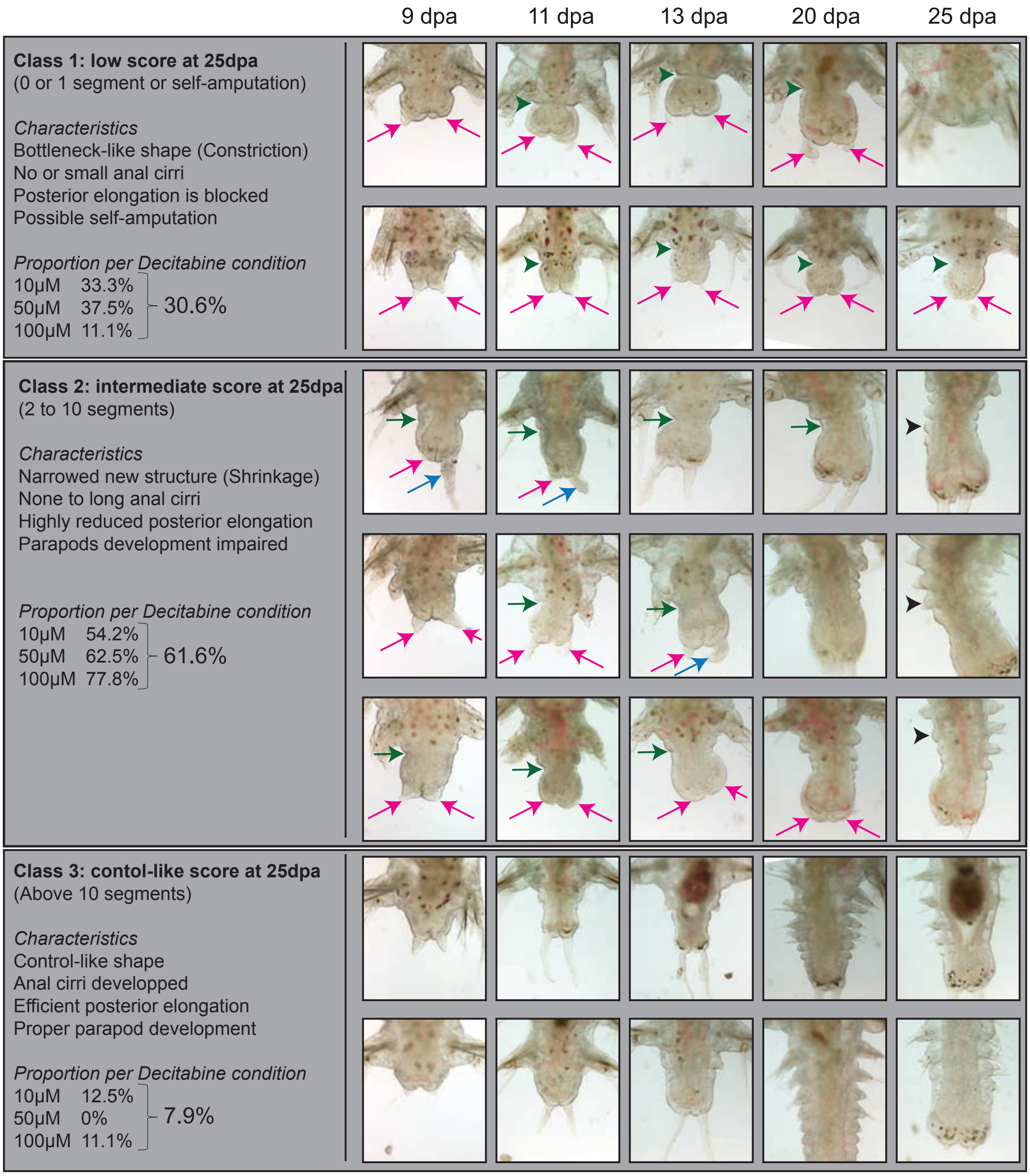

**Additional file 11: Figure S9: Multiple correlation analysis between regeneration and segment addition.** (A-B) Statistical analysis of regeneration score correlation in (A) control and (B) Decitabine treated-worms. Blue dots show positive correlations and red dots are for negative correlations. Dot size is proportional to the correlation score and significant correlations are highlighted with stars. Spearman correlation with Holm post hoc test (* for p <0.05).

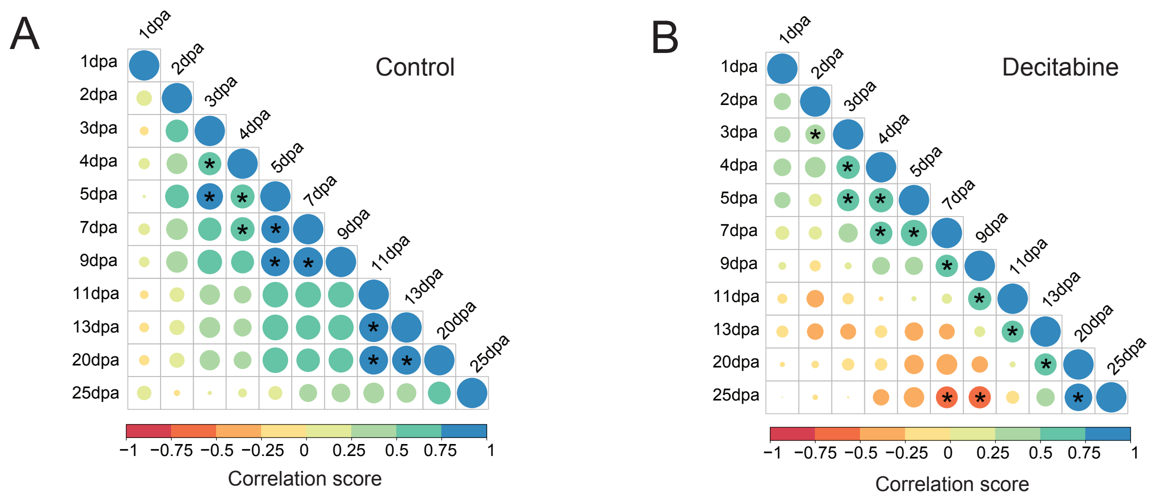

**Additional file 12: Figure S10: Analysis of long-term Decitabine effects on regenerating and growing worms after a second amputation.** (A) Schematic representation of the experimental design. Worms were treated with Decitabine (10µM, 50µM, or 100µM), or DMSO (0,01 %; control) or kept in normal sea water (control) for five days following amputation (5 days post-amputation, dpa). Decitabine was washed out and worms were kept in normal sea water until 25dpa (see Figure 8). A second amputation was performed (which removed the regenerated region) and worms were kept in normal sea water until 18 days post-second amputation (18dpSa). Observations were done at indicated time points (pink bars). (B) Graphic representation of the stages reached by control worms (normal sea water and DMSO 0,01%) and Decitabine-treated worms after the second amputation. A significant delay was observed for worms treated with 10µM as compared to controls at 18dpSa. Worms treated with the other concentrations of Decitabine regenerated and add segments like controls. Two experiments, mean ± SD, 2-way. ANOVA (p-value: Time $p < 0.0001$, Treatment $p = 0.0283$, Interaction $p = 0.0663$) with Tukey post hoc test (*: $p<0.05$). Only p-values corresponding to comparison to normal sea water is shown, as similar to DMSO control comparison.

(C) Most Decitabine worms that were class I or class II at 25dpa (after first amputation) regenerated after a second amputation without any morphological abnormalities (84,2% and 88,1%, respectively), but some of them show minor morphological defects in parapodia and chaetae formation (15,8% and 11,9%, respectively). (D and E) Multiple correlation analysis between regeneration/segment addition after first and second amputation. Only positive correlations were observed for both control and Decitabine-treated worms after a second amputation. Blue dots indicate positive correlations and red dots are for negative correlations. Dot size is proportional to the correlation score and significant correlations are highlighted with stars. Spearman correlation with Holm post hoc test (* for $p <0.05$).

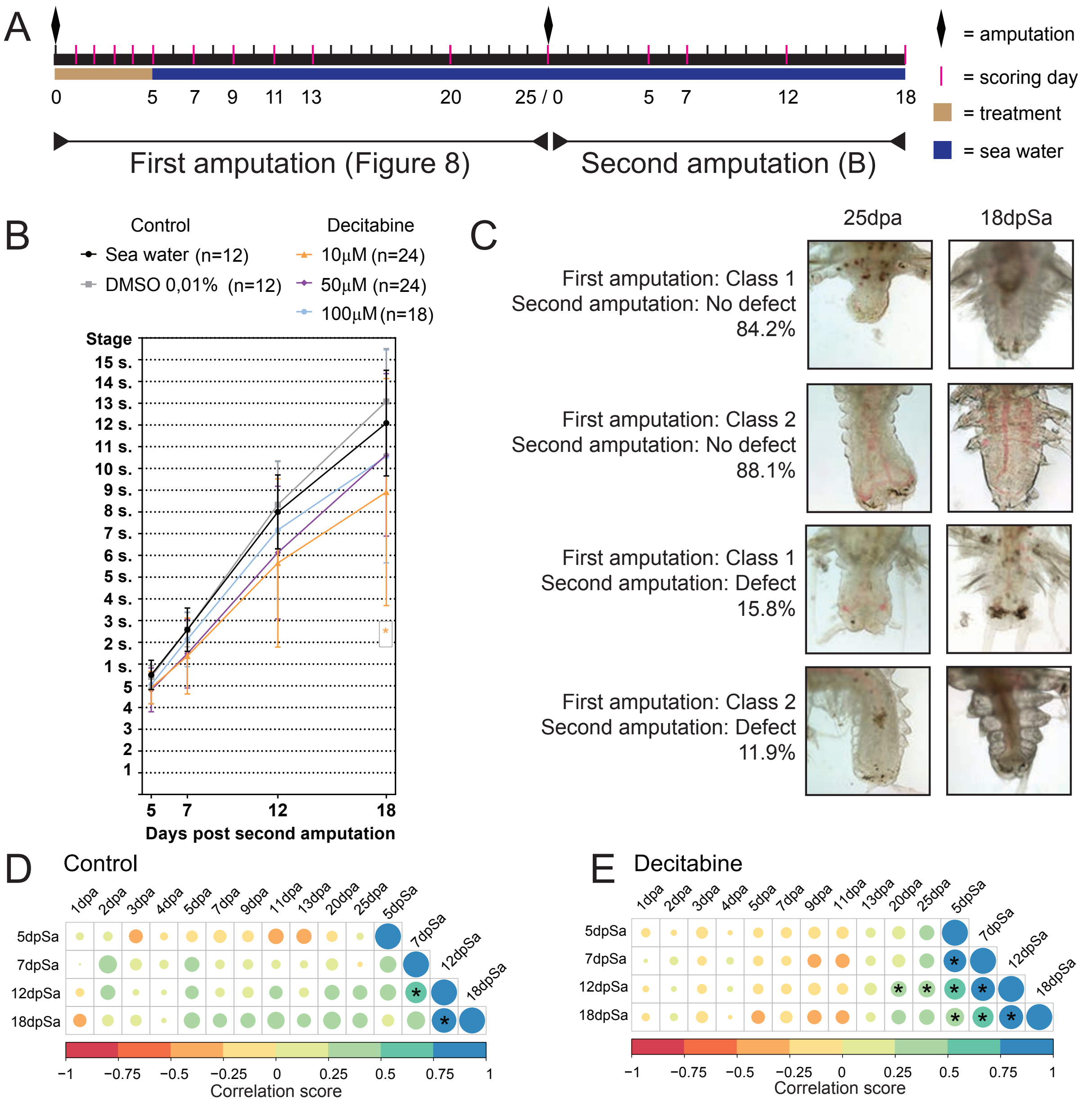
A
= amputation
= scoring day
= treatment
= sea water
0 5 7 9 11 13 20 25 / 0 5 7 12 18
First amputation (Figure 8)
Second amputation (B)
B
Control
Sea water (n=12)
DMSO 0,01% (n=12)
Decitabine
10µM (n=24)
50µM (n=24)
100µM (n=18)
Stage
Days post second amputation
C
25dpa
18dpSa
First amputation: Class 1
Second amputation: No defect
84.2%
First amputation: Class 2
Second amputation: No defect
88.1%
First amputation: Class 1
Second amputation: Defect
15.8%
First amputation: Class 2
Second amputation: Defect
11.9%
D Control
E Decitabine
1dpa 2dpa 3dpa 4dpa 5dpa 7dpa 9dpa 11dpa 13dpa 20dpa 25dpa 5dpSa 7dpSa 12dpSa 18dpSa
Correlation score
